# Genomic Characterization of Rare Earth Binding by *Shewanella oneidensis*

**DOI:** 10.1101/2022.10.31.514631

**Authors:** Sean Medin, Alexa M. Schmitz, Brooke Pian, Kuunemuebari Mini, Matthew C. Reid, Megan Holycross, Esteban Gazel, Mingming Wu, Buz Barstow

**Author notes:** Corresponding author: Buz Barstow, 228 Riley-Robb Hall, Cornell University, Ithaca, NY 14853.

## Abstract

Rare earth elements (REE) are essential ingredients of sustainable energy technologies, but separation of individual REE is one of the hardest problems in chemistry today^1^. Biosorption, where molecules adsorb to the surface of biological materials, offers a sustainable alternative to environmentally harmful solvent extractions currently used for separation of rare earth elements (REE). The REE-biosorption capability of some microorganisms allows for REE separations that, under specialized conditions, are already competitive with solvent extractions^2^, suggesting that genetic engineering could allow it to leapfrog existing technologies. To identify targets for genomic improvement we screened 3,373 mutants from the whole genome knockout collection of the known REE-biosorbing microorganism *Shewanella oneidensis* MR-1^3,4^. We found 130 genes that increased biosorption of the middle REE europium, and 112 that reduced it. We verified biosorption changes from the screen for a mixed solution of three REE (La, Eu, Yb) using Inductively Coupled Plasma Mass Spectrometry (ICP-MS) in solution conditions with a range of ionic strengths and REE concentrations. We found, among other things, that disruptions of a key regulatory component of the arc system (*hptA*), which regulates cellular response to anoxic environments and polysaccharide biosynthesis related genes (*wbpQ, wbnJ, SO_3183*) consistently increase biosorption across all our solution conditions. Our largest total biosorption change comes from our *SO_4685*—a capsular polysaccharide (CPS) synthesis gene—disruption which results in an up to 79% increase in biosorption and *nusA*—a regulatory protein—disruption which results in an up to 35% decrease in biosorption. Knockouts of *glnA, pyrD*, and *SO_3183* increase relative biosorption affinity for ytterbium over lanthanum in multiple solution conditions tested, while many other genes we explored have more complex binding affinity changes. Taken together, these results begin to elucidate how various genes affect the membrane chemistry of *S. oneidensis* and offer potential targets for improving biosorption and separation of REE.

## Introduction

Rare Earth Elements (REE), typically referring to the lanthanides (lanthanum to lutetium) and sometimes scandium and yttrium, are essential ingredients for sustainable energy technologies including high strength lightweight magnets used in electric vehicles and wind turbines^5,6^; room temperature superconductors^7^; lightweight high-strength alloys^8,9^; high-efficiency lighting^10^; and battery anodes^11^. All of these applications put an increasing demand on the global REE supply chain. As the world demand for sustainable energy grows^12^, developing a sustainable supply chain for high-purity REE is critical^13^.

Current methods for refining REE often involve harsh chemicals, high temperatures, high pressures, and generate a considerable amount of toxic waste^14-16^. These processes give sustainable energy technologies reliant on REE a high environmental and carbon footprint.

The majority of REE chemical separations utilize commercially available organic solvents and extractants^17^. All lanthanides exist as trivalent cations and the ionic radius difference between the largest rare earth, La^3+^, and the smallest rare earth, Lu^3+^, is only 0.17 Å^18^. This means that separations of adjacent or near-adjacent REE pose an enormous challenge for conventional chemical methods, requiring organic solvent extractions in extremely long mixer settler devices^19^. This results in large amounts of toxic waste being generated. As a consequence, due to its high environmental standards, the United States has no capacity to produce purified REE. Furthermore, only one REE purification plant exists outside of China^15,16^.

New biological and chemical methods^20-23^ have recently been developed to address the challenges of total^24-30^, light and heavy REE^31^, and individual^1,2,32^ REE-separations.

Biosorption and desorption from the surface of a microbial cell offers an environmentally-friendly route for individual REE-separation. The cell surface, containing proteins^33^, lipids^34^, and polysaccharides^35^, offers a rich chemical environment for selectively binding and releasing REE. The membranes of both gram-negative and gram-positive bacteria contain sites that bind REE^36^. Bonificio *et al*. have already demonstrated nascent REE-separation capability by biosorption and desorption under decreasing pH by *Shewanella oneidensis* MR-1^2^. Furthermore, Bonficio *et al*. demonstrated that, under specialized conditions, REE separations are already competitive with solvent extraction using *Roseobacter* sp. AzwK-3b^2^.

However, biosorption remains poorly understood^25,37,38^. Despite the promising capability identified by Bonificio *et al*., there is no obvious roadmap for genetic engineering of biosorption to reduce binding of non-REE metals and improve binding selectivity for specific REE that would enable separation with biosorption to leapfrog existing technologies. To address this knowledge gap, in this work we comprehensively profile the genetics of REE biosorption in *S. oneidensis*.

## Results

### Genetic Screen Finds 242 Genes that Influence Europium Biosorption

We screened 3,373 unique members of the *S. oneidensis* whole genome knockout collection^3,4^ for genes that control biosorption of the middle REE europium (Eu) using an Arsenazo III colorimetric screen^39^ (**Figures 1A** and **S1, Materials and Methods**). In total, we found 130 gene disruption mutants that have significantly higher Eu-biosorption, and 112 that have significantly lower Eu-biosorption (**Figure 1B, Dataset S1**).

**Figure 1.**
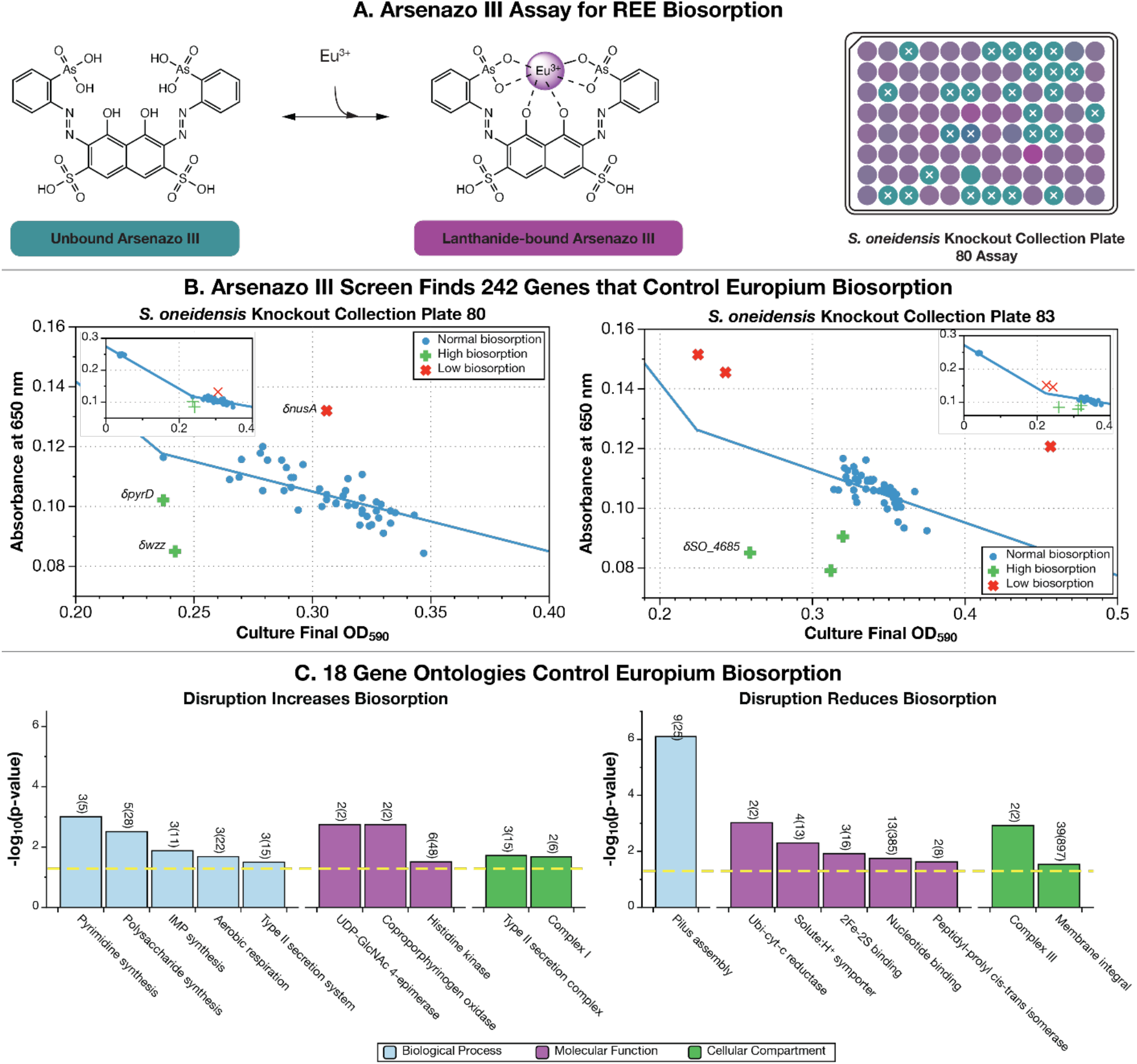
Screening the *Shewanella oneidensis* whole genome knockout collection finds 242 genes representing 18 gene ontologies that control Eu-biosorption. We used the Arsenazo III (As-III) competitive assay for europium- (Eu-) binding [Hoogendorn2018a] to screen 3,373 unique members of the *S. oneidensis* whole genome knockout collection [Baym2016a, Anzai2017a] to identify mutants with modified REE-biosorption capability. (**A**) Unbound As-III absorbance peaks at ≈ 530 nm (resulting in a cyan color), while Eu-bound As-III (proposed structure) absorbance peaks at ≈ 650 nm (purple). Right panel shows a computer-generated image of a sample assay plate derived from spectroscopic data. Higher biosorption by *S. oneidensis* results in a lower concentration of Eu-As-III and hence lower 650 nm absorption (the well will be more cyan-colored) while lower biosorption results in a higher concentration of Eu-As-III (the well will be more purple-colored). Additional information on the high-throughput screen is presented in **Online Methods** and **Figure S1**. (**B**) The As-III screen found 242 genes that control Eu-biosorption (**Dataset S1**). The largest source of Eu-biosorption variability in the screen is due to bacterial density differences between mutants. For most mutants, the optical density of the culture at the start of the biosorption screen will map onto As-III absorption at 650 nm by a linear piecewise function (shown as a blue solid line). Mutants shown as red diagonal crosses had significantly less biosorption than the plate average. Mutants shown as green horizontal crosses had significantly higher biosorption than the plate average (mutants shown as blue dots are not significantly different from the average). (**C**) Gene ontology enrichment analysis found that 18 ontologies were enriched with mutants discovered by the As-III screen. The yellow dotted line indicates a *p*-value of 0.05. We only show results with *p*-values below 0.05 and gene ontologies with > 1 representative mutant. Numbers above each bar indicate the number of significant biosorption genes within each ontology in the screen results relative to the number in the *S. oneidensis* genome. Precise definitions of each gene ontology are shown in **Dataset S2**. IMP: inosine 5′-monophosphate; UDP-GlcNAc 4-epimerase: UDP-N-acetylglucosamine 4-epimerase; Ubi-cyt-c reductase: ubiquinol-cytochrome-c reductase.

### 18 Gene Ontologies are Significantly Enriched Among Genes Influencing REE Biosorption

Using Fisher’s exact test, we analyzed each set of gene disruption mutants—those with higher or lower Eu-biosorption—for enrichment of gene ontologies to identify trends in the overall set of genes contributing to biosorption. We identified 18 gene ontologies that were significantly enriched (*p* < .05) and had more than one representative gene within our genetic screen results (**Figure 1C, Dataset S2, Materials and Methods**). Ten were enriched among genes whose disruption increases biosorption while eight were enriched among genes whose disruption decreases biosorption. Ontologies discussed in this work whose gene disruptions increase Eu-biosorption include pyrimidine synthesis, polysaccharide synthesis, and histidine kinase activity ontologies, while one ontology discussed in this work whose gene disruptions decrease Eu-biosorption is pilus assembly.

### 13 Operons are Significantly Enriched in Genes Influencing REE Biosorption

Using computationally produced predictions for operons in *S. oneidensis*^40,41^ and Fisher’s exact test, we identified thirteen operons with statistically significant (*p* < 0.05) enrichments of genes whose disruptions produced differential biosorption in the Arsenazo-III screen (**Figure S2, Dataset S3, Materials and Methods**). Out of the thirteen, three operons had highly significant enrichment of hits (*p* < .001). Identifying enriched operons allowed us to further refine our analysis of which functions contribute to biosorption. For example, while ‘polysaccharide synthesis’ is highlighted as relevant from our ontology analysis, our operon analysis points us to two specific polysaccharide and O-antigen synthesis operons (‘PS1’ and ‘PS2’) that impact biosorption (**Figures 2A** and **B**). Our operon analysis also specifies the MSHA pilus—previously highlighted more generally in the ‘pilus assembly’ ontology in our ontology enrichment analysis—as important to REE biosorption as 12 out of the 15 gene disruptions in this operon produced lower Eu-biosorption in our screen. Many, but not all, of the genes identified in these three operons are also highlighted in the corresponding ontologies.

**Figure 2.**
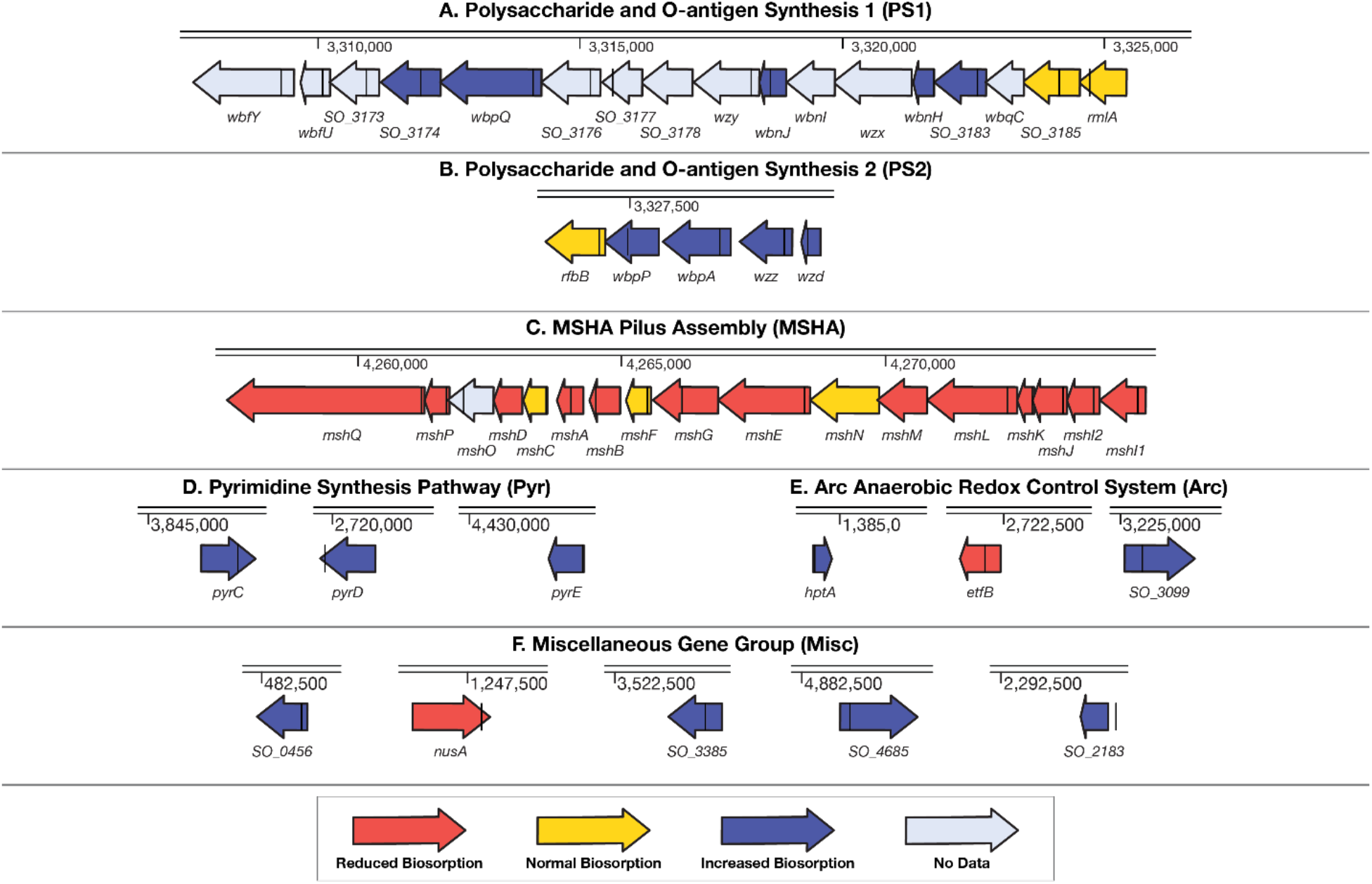
Operon enrichment, ontology enrichment, and regulatory analyses pinpoint 6 groups of genes that influence multiple mechanisms behind Eu-biosorption by *S. oneidensis*. (**A** to **C**) The results of the high-throughput Eu-biosorption screen (**Dataset S1**) of the *S. oneidensis* knockout collection [Baym2016a, Anzai2017a] were analyzed to find operons with statistically-significant enrichments of hits (**Dataset S3**). The location of the transposon disruption in each gene is marked as a black line. Here we show three operons that are the most statistically-significant results of this analysis. (**D**) The pyrimidine synthesis pathway was selected by ontology enrichment analysis (**Dataset S2**). (**E**) One gene involved in the Anaerobic Redox Control (Arc) regulatory system (*hptA*), as well as two genes regulated by Arc whose knockouts produced differential biosorption were also selected for further analysis. (**F**) Finally, five genes whose knockouts produced some of the largest changes to Eu-biosorption were also selected for further analysis.

### Regulatory Analysis of the Arc System Highlights Several Other Genes Important to Biosorption

A disruption in the histidine kinase *hptA* was found to substantially increase biosorption in our genetic screen and *hptA* contributes to the enrichment of the ‘histidine kinase’ ontology group among hits. Since HptA is a regulatory protein, we speculated that its impact on biosorption must be caused by indirect activation or repression of downstream genes and thus sought to pinpoint the genes responsible.

HptA is part of the two-component Anoxic Redox Control (Arc) system in *S. oneidensis*. ArcS phosphorylates HptA in the absence of O_2_, and HptA in turn phosphorylates the response regulator ArcA^42^. While we did not have gene disruptions we could screen for *arcS* or *arcA* it is reasonable to assume that disruption of *hptA* produces a similar effect as disruption of *arcS*. We found that individual disruption of 29 of the 604 genes whose activity is affected by an *arcS* deletion (and thus likely a *hptA* deletion as well)^42^ produced significant changes in Eu-biosorption (**Table S1**).

Of the ArcS-regulated genes that may contribute to biosorption, several disruptions decreased biosorption (such as δ*etfB*, a disruption of subunit B of the *e*^-^ transfer flavoprotein^43^); several increased biosorption (such as δS*O_3099*, a disruption of an outer membrane long-chain fatty acid receptor^43^); and several were also found to contribute to enriched ontologies (such as *δpyrE* which is involved in pyrimidine biosynthesis^43^). (‘δ’ indicates a gene disruption mutant and ‘Δ’ indicates a gene deletion mutant.)

### Six Groups of Genes that Influence Multiple Mechanisms of REE Biosorption were Chosen for Detailed Analysis

Our As-III biosorption screen was limited because it could analyze only a single REE. We thus sought to expand our analysis by looking at how our gene disruptions impacted biosorption of multiple REE. We selected six groups of genes representing a wide range of cellular functions for detailed analysis with the Inductively Coupled Plasma Mass Spectrometry (ICP-MS)—an instrument for making robust measurements of concentrations of multiple elements—in order to validate the results we found with the As-III assay. We selected these groups based on gene ontology enrichment, operon enrichment, and regulatory analyses from our Eu-screen biosorption data.

For three of these groups, we selected operons of interest: polysaccharide and O-antigen synthesis operons 1 and 2 (PS1 and PS2; **Figures 2A** and **B**) and the MSHA pilus operon (MSHA; **Figure 2C**). Within the MSHA operon, we chose to look at δ*mshQ*, δ*mshD*, δ*mshC*, δ*mshA*, δ*mshB*, δ*mshL*, δ*mshJ* because these are all predicted to be either outer membrane proteins or found on the pilus appendages^44^. All of these, except for δ*mshC*, had significantly lower biosorption in the As-III Eu-biosorption screen.

An additional group of disruptions in non-contiguous genes (δ*pyrC*, δ*pyrD*, and δ*pyrE*) was selected for detailed study based on their contribution to the enrichment of the pyrimidine biosynthesis gene ontology (Pyr; **Figure 2D**).

In addition to genes contributing to ontology and operon enrichment, we chose to analyze disruptions in genes from the Arc system, including a disruption of the Arc system histidine kinase *δhptA* as well as disruptions of two genes regulated by the Arc system: *δetfB* and *δSO_3099* (Arc; **Figure 2E**).

As a miscellaneous group, we selected five gene disruption mutants that were independent of any identified grouping but produced strong changes in biosorption (Misc; **Figure 2F**). A mutant in the gene coding for the transcriptional regulator NusA (δ*nusA*)^43^ (also see **Figure 1B**, left panel) had lower biosorption. Meanwhile disruption of *SO_4685*, which codes for a protein involved in extracellular capsular polysaccharide synthesis (CPS)^43^ produced higher biosorption (also see **Figure 1B**, right panel). Likewise, disruption of *SO_3385* which codes for a transcriptional activator of singlet oxygen protection^43^ produced higher biosorption. Finally, an insertion 150 bp upstream of *SO_2183*, which codes for a protein involved in biosynthesis of the cell wall component peptidoglycan^43^, had higher biosorption.

### ICP-MS Validates Differential Biosorption of Genes from Selected Gene Groups

ICP-MS measurements of mixed REE-biosorption largely validated the results of the high-throughput biosorption assay (**Figures 3A** to **D, Table S2**). We measured biosorption of lanthanum (La; representing light REE), Eu (for middle REE), and ytterbium (Yb; for heavy REE) under four solution conditions (**Materials and Methods**) by 25 gene disruption mutants representing the six groups of genes chosen for detailed analysis (**Figures 3A** to **D**) and six clean deletion mutants (**Figures 3E** to **H**).

**Figure 3.**
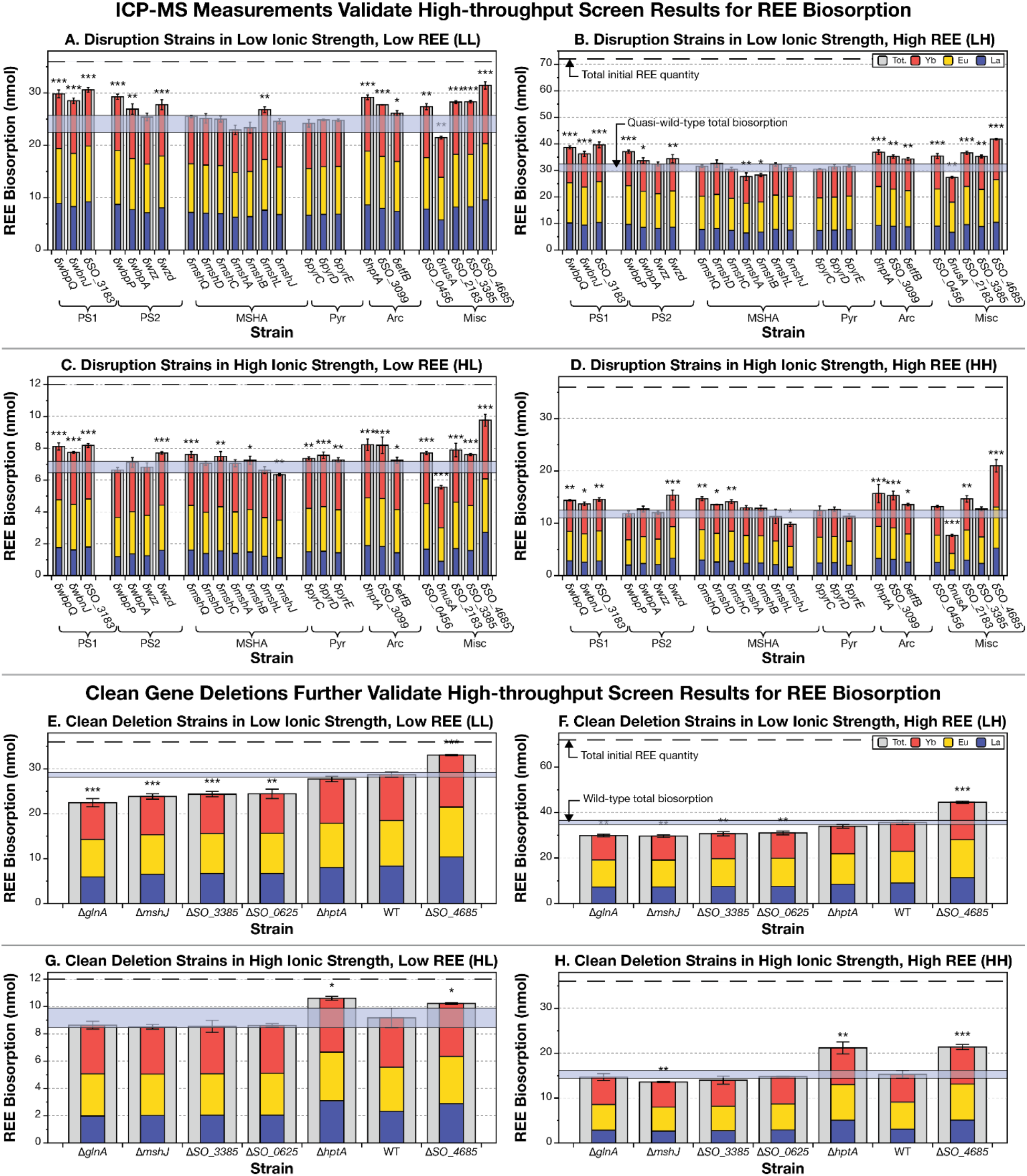
ICP-MS measurements validate the results of high-throughput Eu-biosorption screening in up to 79% of cases. Bar plots show levels of lanthanum (blue), europium (yellow), ytterbium (red), and total REE (grey) biosorption for each strain. The error bar indicates the standard deviation on the total biosorption of three biological replicates. The number of stars above each bar indicates the statistical significance (two-tailed t-test) of the measurement difference from quasi-wild-type (**A** to **D**) and wild-type (**E** to **H**): *: *p*-value < 0.05; **: *p*-value < 0.01; ***: *p*-value < 0.001. δ indicates a transposon insertion mutant (in panels **A** to **D**), while Δ indicates a clean deletion mutant (in panels **E** to **H**). Cross-checks of As-III Eu-biosorption assay and ICP-MS measurements with transposon mutants are shown in **Table S2**. (**A**) The low ionic strength, low initial REE concentration environment matches 53% of the As-III screen **(Table S2). (B)** The low ionic strength, high initial REE concentration environment (LH) recapitulates the highest percentage (63%) of results of the As-III screen. (**C**) The high ionic strength, low REE environment (HL) reproduces 63% of significant changes to biosorption. (**D**) The high ionic strength, high initial REE environment reproduces the smallest number (42%) of results from the As-III screen. (**E** to **H**). Clean deletion mutants replicated at least some of the results of transposon mutant measurements in three of four cases.

In industrial settings, REE are processed in a wide array of combinations and concentrations, and with a wide range of competing metal concentrations. We chose to explore two axes of interest to characterize our selected insertion mutants: low and high ionic strength (provided by sodium chloride); and (2) low and high total initial REE concentration. We speculated that testing different ionic strength concentrations would be of particular interest because of the possibility of cation competition for REE binding sites and the ability of ionic strength to change protein configurations among other potential impacts. In short, we have four solution conditions: low ionic, low initial REE concentration (LL); low ionic, high REE (LH); high ionic, low REE (HL); and high ionic, high REE (HH). Biosorption solution conditions used are detailed in **Table 1**.

**Table 1.** Solution environments for detailed REE biosorption measurements. All solutions were adjusted to pH 5.5.

| Environment | Media Composition | Initial REE Concentration | Initial Total REE Quantity (nmol) |
| --- | --- | --- | --- |
| LL (Low Ionic, Low Initial REE) | 10 mM MES + 10 mM NaCl | 30 μM each La, Eu, Yb | 36 |
| LH (Low Ionic, High Initial REE) | 10 mM MES + 10 mM NaCl | 60 μM each La, Eu, Yb | 72 |
| HL (High Ionic, Low Initial REE) | 10 mM MES + 50 mM NaCl | 10 μM each La, Eu, Yb | 12 |
| HH (High Ionic, High Initial REE) | 10 mM MES + 50 mM NaCl | 30 μM each La, Eu, Yb | 36 |

As a benchmark for comparison, we selected 4 quasi-wild-type (qWT) transposon insertion mutants from the *S. oneidensis* whole genome knockout collection that had the insertion in a location unlikely to impact biosorption (**Materials and Methods**). We found that the genuine wild-type *S. oneidensis* showed at least 13% higher biosorption than the average qWT in every solution condition (**Figure S4**), indicating that the presence of a transposon insertion alone may affect biosorption. We thus chose to compare our notable transposon mutants with the average of our qWT mutants rather than the true wild-type.

Our multiple-REE biosorption assay recapitulated significant increases or decreases in biosorption from the As-III screen for 54% (13/24) of disruption mutants tested for the LL environment (**Figure 3A**), 62% (15/24) for the LH (**Figure 3B**) and HL (**Figure 3C**) environments, and 42% (10/24) for the HH (**Figure D**) environment. 79% (19/24 genes) of our insertions are validated in at least one environment (**Figure 3** and **Table S2**).

Disruptions to 9 genes produced higher biosorption in all environments: disruption of the uncharacterized protein Wzd (δ*wzd*) (Polysaccharide Synthesis Operon 1), disruption mutants for all 3 of the chosen Polysaccharide Synthesis Operon 1 genes, disruptions to all 3 genes related to the Arc system (including δ*etfB*, which had lower biosorption in the genetic screen), and disruption of the LD-transpeptidase encoding gene *SO_2183* and the CPS-synthesis gene *SO_4685* from the Miscellaneous group. In fact, δ*SO_4685* produced our largest observed increases in total biosorption ranging between 31% (in LL) and 79% (in HH) higher than the qWT.

The insertions for *SO_0456* and *SO_3385* (both in the Miscellaneous group) had significantly higher biosorption than the qWT in every solution condition except for HH.

Only one gene disruption produced consistently lower total REE-biosorption. Specifically, disruption of the last 10% of the coding region for the transcription regulator NusA (δ*nusA*; the Miscellaneous group), produced our largest observed reductions in total biosorption ranging between 11% (in LL) and 35% (in HH) lower than the qWT.

Five gene disruption mutants showed a notable discrepancy in total biosorption between low ionic strength (**Figures 3A, B, E**, and **F**) and high ionic strength (**Figures 3C, D, G**, and **H**) environments. Most notably, the MSHA genes showed the greatest environment dependency in their biosorption results. For example, δ*mshJ* had significantly lower biosorption in the high ionic strength cases (HL and HH; **Figures 3C** and **D**), but no significant change in the low ionic strength cases. Meanwhile, δ*mshA* and δ*mshB* (MSHA Pilus had lower biosorption capabilities in the low ionic strength environments (LL and LH; **Figures 3A** and **B**), and either no significant change or a biosorption increase in the high ionic strength environments.

All three pyrimidine biosynthesis gene insertions had significantly higher biosorption (between 7 and 11% higher than the qWT) in the HL environment but registered no significant difference in any other environment (**Figure 3D**).

### Clean Gene Deletion Mutants Largely Verify Biosorption Results for Gene Disruption Mutants

We created clean deletion mutants for four genes whose disruption conferred standout biosorption changes: Δ*mshJ*, Δ*hptA*, Δ*SO_3385*, Δ*SO_4685*. Three of four of these clean deletion mutants at least partially reproduced the results of the corresponding disruption mutants. While insertion mutants are effective at knocking out gene function, they do not always successfully mimic a true single gene knockout. While the *S. oneidensis* knockout collection was designed to mitigate polar effects^45-47^, it is possible that the early parts of the knockout gene can still create a partially functional product^48^, which could account for the discrepancies between deletion and disruption strains. We also created two additional mutants for genes identified by the Arsenazo-III genome-wide screen (**Dataset S1**): Δ*SO_0625* (a knockout for periplasmic cyctochrome c oxidase regulatory protein) and Δ*glnA* (for glutamine synthetase).

Clean deletion of the pilus biogenesis gene *mshJ* (Δ*mshJ*) had significantly lower biosorption (11 to 17%) in three of the four environments tested (all except HL where the biosorption level was lower, just not statistically significantly). The main difference between the insertion and the clean deletion mutant is that Δ*mshJ* produces lower biosorption in the low ionic strength environments, while δ*mshJ* has no significant change compared to the qWT in those environments.

Clean deletion of the transcriptional regulator gene *hptA* (Δ*hptA*) produced large increases in biosorption (16 to 39%) in high ionic strength environments (HL and HH; **Figures 3G** and **3H**) just like the transposon mutant (δ*hptA*) (**Dataset S1, Figures 3C** and 3**D**). However, unlike the transposon mutant, Δ*hptA* did not produce significantly different biosorption from the wild-type in low ionic strength environments (LL and LH; **Figures 3E** and **F**).

Complete knockout of the capsular EPS biosynthesis gene SO_4685 (Δ*SO_4685*) showed a significant increase in biosorption (15 to 40%) in every environment just like the insertion mutant (**Figure 3**).

Clean deletion mutants for *SO_0625* and *glnA* produced significantly lower biosorption than the wild-type in the low ionic strength cases, and non-statistically significantly lower biosorption in the high ionic strength cases.

### Nine Insertion Mutants Have Notable Modification of Individual Lanthanide Binding Preference

We next examined if, in addition to changes to total REE biosorption, our mutants produced changes to the relative biosorption affinity for particular REE over others. Out of the 25 insertion mutants we conducted follow up ICP-MS analysis on, nine of our mutants appeared to produce robust changes to biosorption preferences for particular REE.

Notably, our qWT already has a marked preference for heavier REE. This preference for heavier REE increases with ionic strength. From an initially equimolar mixed REE solution of La, Eu, and Yb, the qWT-biosorbed fraction contains between ≈ 19% and ≈ 28% La; ≈ 37% and 40% Eu; and ≈ 35% and 44% Yb (**Figure S4**).

For most transposon insertion mutants, under most of the four solution conditions tested, individual REE biosorption is linearly related to total biosorption (**Figure 4, Materials and Methods**) over a finite range (note the finite extent of dashed black lines in **Figures 4A** to **B**).

**Figure 4.**
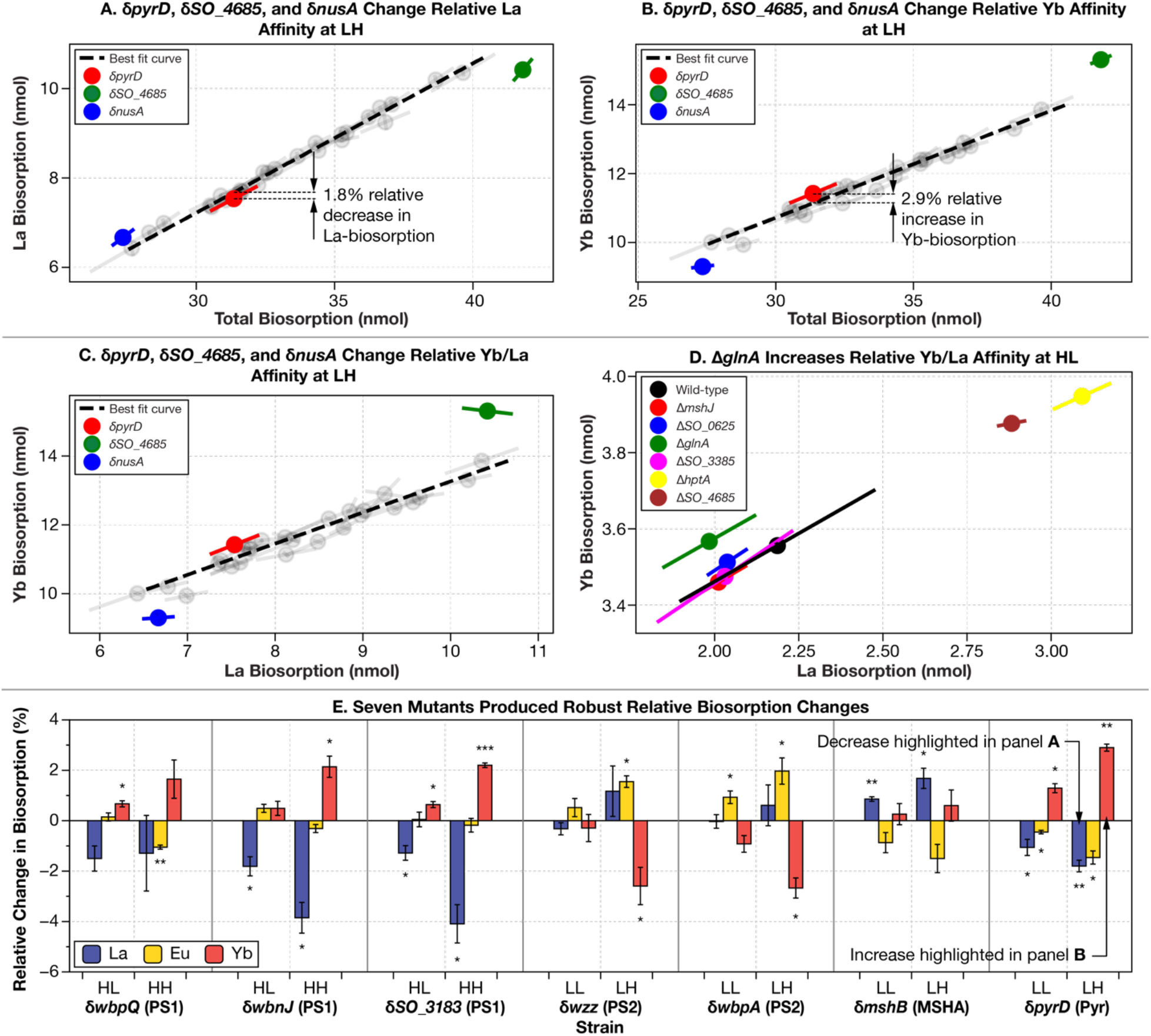
Nine gene disruption mutants make notable changes to REE-biosorption selectivity. (**A** to **C**) For most transposon insertion mutants (including those with modified total REE biosorption), under most of the four solution conditions tested, individual REE biosorption is linearly related to total biosorption or individual biosorption of either of the other 2 REE tested (grey circles, and the black dashed fit lines in panels **A** to **C**) over a finite range of REE biosorption (note the finite extend of dashed black lines in panels **A** to **C**). (**A** to **D**) Individual points indicate the mean values of the mutants and the error bars show the standard deviation of three biological replicates along the axis of maximal variation. (**A** to **C**) We highlight changes in La and Yb affinity for δ*pyrD* in the LH environment as well as two mutants (δ*nusA* and δ*SO_4685*) who’s total biosorption was too small (δ*nusA*) or large (δ*SO_4685*) to compare to our finite line of best fit, yet nonetheless clearly had different La and Yb affinity relative to the other mutants. In particular, note how both δ*nusA* and δ*SO_4685* have similar La biosorption to other mutants in (**C**) yet had very different Yb biosorption. (**D**) We had insufficient data to perform a line of best fit on our clean gene deletion data, yet it is clear from this plot that the *glnA* deletion has an increase in relative Yb/La affinity. (**E**) Here, we display all our mutants with robust biosorption changes (mutants that enhanced or decreased relative biosorption in multiple environments, or when a particular REE had enhanced or lowered relative biosorption in more than half the genes in a particular group of interest). The number of stars above or below each bar indicates the statistical significance (two-tailed t-test) of the measurement difference from quasi-wild-type: *: *p*-value < 0.05; **: *p*-value < 0.01; ***: *p*-value < 0.001. δ indicates a transposon insertion mutant, Δ indicates a clean deletion mutant. Error bars indicate standard deviation of three biological replicates. PS1 and 2: Polysaccharide Synthesis 1 and 2; MSHA: MSHA Pilus Assembly; Pyr: Pyrimidine Synthesis.

However, every transposon insertion mutant that we tested had at least one REE biosorption result in at least one environment that deviated from the linear individual-to-total relationships established for most mutants under most conditions (**Figures 4A** to **4C**, and **S4**). It is notable, however, that having used a p-value of only .05 while performing 12 significance tests per mutant, our statistical test was not very stringent. We thus narrowed our criteria to select for ‘robust’ results by focusing on only those mutants with enhanced (or decreased) relative biosorption of a particular REE in the same direction in multiple environments or when a particular REE had enhanced or lowered relative biosorption (again, in the same direction) in more than half the genes in a particular group of interest.

Seven mutants produced robust results according to our criteria. These highlighted mutants are summarized in **Figure 4E**.

Disruption of genes in the Polysaccharide Synthesis 1 operon tend to increase Yb-binding and decrease La-binding for high ionic strength environments (**Figure 4E, Figure S6**). For example, disruption of *SO_3183* increases relative Yb-binding under HH by 2.2% and reduces relative La-binding by 4.1%.

Disruption of two of the four genes in the Polysaccharide Synthesis 2 operon (*wzz* and *wbpA*) produces a significant increase in Eu and a significant decrease in Yb biosorption in the LH condition (**Figure S6**). The disruption of *wbpA* has a significant relative increase in binding of Eu under the LL condition as well.

Disruption of *mshB* (δ*mshB*) produces significant reductions of in La-binding in low ionic strength conditions, likely at the expense of Eu-binding (although these changes are not significant).

Disruption of *pyrD* produces significant increases in Yb-binding coupled to reductions in La- and Eu-binding in both LL and LH. Under LH, the increase in Yb-binding of 2.9% is one of the largest significant increases in relative REE binding.

While disruption of *pyrC* under HH produces the largest significant change, increasing La-binding by 5.8%, it did not meet our robustness metric.

Two insertion mutants from the Miscellaneous group (δ*nusA* and δ*SO_4685*) have total biosorption levels that are so different from the rest of our insertion mutants that we did not include them in our formal analysis of relative REE changes (they were outside of the finite linear fit region in **Figure 4A-C**). However, in the case of the LH environment, we still found a clear way of illuminating relative REE affinity changes (**Figure 4C**). δ*nusA* produces very similar La-binding to other transposon insertion mutants that were in-range for our analysis. At the same time, it had a much lower relative level of Yb biosorption. This made it clear that δ*nusA* had relatively higher La-binding and relatively lower Yb-binding. Similarly, δ*SO_4685* had a similar level of La-binding compared to other in-range strains, but bound a much greater amount of Yb, implying relative increase in preference for heavy versus light REE.

## Discussion

Our genetic screen for biosorption reveals layers of the outer surfaces (inner and outer membranes, and periplasmic layer) of *S. oneidensis* that modulate access to REE-binding sites in *S. oneidensis*. These layers include polysaccharides (synthesized by Polysaccharide Synthesis Operons 1 and 2), MSHA pili (synthesized and assembled by the MSHA Pilus Assembly Operon), and a variety of outer membrane proteins (SO_0456, SO_3099, MshQ, MshL, MshJ).

### Disruption of Polysaccharide Synthesis Operon 1 Raises Biosorption

We speculate that disruption of Polysaccharide Synthesis Operon 1 (PS1) modifies the lipopolysaccharide (LPS) layer on the outer membrane of *S. oneidensis*. Many of the genes coded by PS1 are responsible for the synthesis of O-antigens, a major component of the LPS^49^. Disruption of all genes selected for further analysis in PS1 increase total biosorption under all solution conditions (**Figure 3**) and generally increase relative Yb-binding and decrease relative La-binding in high ionic strength conditions (**Figure 4E**). Among the three genes tested in the PS1 group, only δ*SO_3183* is directly implicated in polysaccharide synthesis (it was highlighted in our gene ontology analysis), although the similar biosorption effects of each of our three mutants seem to suggest that they may all be part of a single pathway.

### Disruption of Polysaccharide Synthesis Operon 2 Modifies the Cell Membrane and REE Biosorption

The disruption mutants selected for in-depth analysis belonging to Polysaccharide Synthesis Operon 2 (PS2) generally cause significant increases in REE-biosorption in at least some cases, although the results were not necessarily consistent from gene to gene. For example, while δ*wbpP* and δ*wbpA* significantly raise biosorption only in the low ionic strength cases, δ*wzd* increases biosorption in every case and δ*wzz* fails to significantly alter total biosorption at all. We suspect this is due to each of the genes in this group having a unique impact on biosorption.

We speculate that disruption of WbpP (δ*wbpP*) raises REE-biosorption in low ionic strength conditions due to its significant role in membrane composition. WbpP transforms UDP-N-acetyl-D-glucosamine to the UDP-N-acetyl-D-galactosamine^50^. In *P. aeruginosa*, WbpP plays a role in the synthesis of B-band O-antigens, a component of the lipopolysaccharide layer^51^. In *V. vulnificus*, deletion of *wbpP* causes the failure of CPS (capsular polysaccharide) formation^52^. The deletion also results in increased cell aggregation, hydrophobicity, and adherence to abiotic surfaces, all suggestive of substantial membrane changes^52^. Without more *S. oneidensis* specific data, it is impossible to know the exact mechanism for δ*wbpP* having increased biosorption in only the low ionic strength cases. However, one possibility could be that, like in *P. aeruginosa*, B-band O-antigens are deleted. Consequently, binding sites that are normally covered by those O-antigens could be revealed that bind to REE only in low ionic strength environments.

Likewise, we speculate that disruption of *wbpA* (δ*wbpA*) also raises biosorption in low ionic strength conditions due to its role in membrane composition. WbpA, like WbpP, takes UDP-N-acetyl-D-glucosamine as a substrate (but transforms it to uronic acid^50^ instead), and is thought to be a key protein in O-antigen biosynthesis in *P. aeruginosa*^53^.

### MSHA Genes Have Highly Environmentally Dependent Effects on Biosorption

The importance of the interaction of the solution environment with gene disruption on biosorption is most strongly illustrated by disruptions of the MSHA Pilus Assembly Operon (MSHA) genes. While 6/7 of the gene disruptions tested (all except δ*mshC*) had lower biosorption in the original screen, only 3/7 had significantly lower biosorption in any of the solution conditions selected for follow up testes and none of them had significantly lower biosorption for every solution condition.

MshA is responsible for forming the main subunit of the pilus and knocking it out thus has a major impact on the MSHA pilus^54^. δ*mshA* had significantly lower biosorption in the low ionic strength conditions. This seems to suggest that the MSHA pilus plays an important role in binding to REE in low ionic strength cases.

At the same time, δ*mshA* has no significant change in biosorption in the high ionic strength conditions, suggesting that the high NaCl concentration is preventing REE from binding to the pili.

### Disruption of Pyrimidine Synthesis Group Increases REE-Biosorption Under High Ionic Strength, Low REE Conditions

Disruption mutants of the Pyrimidine Synthesis Group genes *pyrC, pyrD*, and *pyrE* all increase biosorption in the high ionic strength, low REE condition. These genes form a section of a pathway in pyrimidine metabolism that produces Orotidine 5′-monophosphate (OMP) from carbamoyl aspartate. The exact mechanism through which these genes effect biosorption remains uncertain and require further investigation.

### Down-regulation of Fatty Acid Transport Regulated by the Arc System is Likely an Important Contributor to Biosorption Changes Observed in δ*hptA*

The large number of biosorption-affecting disruptions in genes regulated by the Arc system suggests that biosorption changes caused by the *hptA* insertion and deletion are the aggregate result of changes to many REE-binding sites in *S. oneidensis*.

Disruption to the gene encoding *SO_3099* causes a substantial and significant increase in biosorption in every solution condition tested. Given that disabling of the Arc regulatory system sensing mechanism results in near elimination of gene expression of *SO_3099*^42^ (**Table S1**), it seems likely that *SO_3099* is one of the major—although likely by no means the only—contributors to the biosorption changes observed in δ*hptA. SO_3099* codes for an outer membrane β-barrel protein that is homologous to the long-chain outer membrane fatty acid transporter FadL in *E. coli*^55^. We speculate that removal of fatty acid transport to the outer membrane of *S. oneidensis* either creates opportunities for increased copy number of other REE-binding compounds on the membrane or increases the accessibility of existing sites.

### Disruptions to *nusA, SO_4685*, and *SO_2183* Have Large Effects on Biosorption

Disruption of *nusA* produces our largest observed reduction in total biosorption (**Figure 3**) and one of the largest relative reductions in Yb-biosorption (at least in the LH case, **Figure 4B**). The disruption occurs in the last 10% of *nusA* which codes for a putatively essential transcription termination/anti-termination protein NusA^56^. This result suggests that the final 10% of *nusA* encodes a domain that plays an important role in REE biosorption, but that is unrelated to NusA’s core essential function in transcription.

δ*SO_4685* produced the highest biosorption of any transposon strain in every environment. Furthermore, δ*SO_4685* produces a large increase in relative Yb-binding. We speculate that disruption of SO_4685, a protein involved in capsular polysaccharide synthesis, improves the accessibility of REE-binding sites on the surface of *S. oneidensis*.

*δSO_2183* produces significantly higher biosorption in every environment tested. We speculate that δ*SO_2183* somehow increases accessibility to REE binding sites on the membrane. *SO_2183* encodes a LD-transpeptidase involved in cross-linking of the peptidoglycan layer that is sandwiched between the inner and outer membranes of *S. oneidensis*^57^. δ*SO_2183* contains a transposon insertion 150 bp upstream of the coding sequence of *SO_2183*, likely in a promoter region.

### Binding Site Changes from Single Gene Knockouts Tend to Have Multiple Effects

While a handful of gene disruption mutants we looked at had consistent biosorption changes across every condition tested, for many of our other genes, changes in biosorption levels were inconsistent across different solution conditions. Some of these discrepancies have simple explanations. For example, several gene insertion mutants (such as polysaccharide synthesis protein δ*wbpP*) had higher biosorption for low ionic strength, but similar biosorption to the qWT for high ionic strength. In the case of δ*wbpP*, it is possible that this is because elimination of *wbpP* results in increased accessibility to binding sites that are capable of binding to REE in low ionic strength cases but not high ionic strength cases—possibly due to competition with sodium ions for binding sites. Thus, knocking out *wbpP* increases REE biosorption in the low ionic strength cases, but has no effect in the high ionic strength cases.

Some other features of our data necessitate more complicated explanations. Most prominently, not a single strain had statistically significant and consistent changes in relative affinity for individual REE across all environments.

Intuitively, one might expect that most gene insertion/deletions that affect biosorption would affect a single binding site. That the gene would encode a single outer membrane protein or a protein that alters some single REE binding lipid or polysaccharide found on the membrane. Based on our genetic screen results, this appears to rarely be the case. More often, gene knockouts likely cause a cascade of effects on other genes resulting in changes to multiple binding sites. Gene disruptions can also have confounding effects unrelated to REE binding site composition. It is possible that some gene disruptions impact the shape of the bacteria or perhaps the optical density to bacteria ratio (since the optical density is what we use to normalize the bacterial density from assay to assay). Perhaps some of these gene disruptions affect secretions of *S. oneidensis* that might interfere with biosorption by competing with surface binding sites^36^. Thus, while our work may answer the question of what genes are important to biosorption, the reasons why they are important almost uniformly requires further investigation.

## Conclusions

We have conducted one of the most comprehensive screens of the genetics of REE-biosorption (or any element for that matter) to date. Our dataset of genes involved in Eu-biosorption gives us and fellow investigators an extensive catalog of sites for further investigation of REE-biosorption.

At the outset of this work, we anticipated that the genetic screen of REE-biosorption would identify a suite of genes encoding individual REE-binding sites on the surface of *S. oneidensis*—either in the form of outer membrane proteins or proteins that create compounds found on the outer membrane. The reality is much more complex.

Many of the gene disruptions that affect total REE-binding and modify the REE-binding preference of *S. oneidensis* likely have functions that affect the outer surface. Both Polysaccharide Synthesis Operon 1 and 2 are involved in making O-antigens on the lipopolysaccharide layer. The MSHA genes synthesize pili on the outer membrane of *S. oneidensis*. Finally, two members of the Miscellaneous Group are related to other outer membrane structures: SO_2183 is involved in synthesis of the peptidoglycan layer and the SO_4685 protein is involved in synthesis of capsular polysaccharides.

Two individual gene disruptions stood out in their impact on biosorption. In every solution condition, disrupting *SO_4685* resulted in the highest biosorption observed, while disrupting *nusA* resulted in the lowest biosorption observed.

Despite there being few if any dominant players in REE-biosorption in *S. oneidensis*, our genetic screen results still provide a roadmap for creating strains of *S. oneidensis* with improved separation of individual REE.

Given that significant headway has been made in separation of lanthanides from competing metals typically found with REE^24,26-29^, we have the potential to control what kind of biosorption environment we have when we mix REE with our engineered bugs.

We thus can use our knowledge of genes that increase relative Yb to La binding to engineer an improved microbe for separating heavy from light REE. We would target genes that are effective at increasing the relative Yb affinity (usually at the expense of La) in a variety of environments. Specifically, we would simultaneously knock out genes whose disruptions improved separations in the low ionic strength cases (*SO_4685* and *pyrD*) as well as genes that improved separation in the high ionic strength cases (*glnA* and the PS1 genes).

Although harder, we can also attempt to build a strain with an increased relative preference for europium. For example, we can simultaneously delete both *wbpA* and *wzz* and see if the resulting microbe has an improved preference for europium.

This will allow us to engineer bacterial cells with greater affinity for particular Rare Earth Elements in order to produce a new, environmentally benign method of extracting and separating REE from other elements and each other.

## End Notes

### Data Availability

The datasets generated during and analyzed during the current study are available from the corresponding author (B.B.) on reasonable request. Datasets for figures are available on Cornell eCommons.

### Code Availability

The Knockout Sudoku software is available at https://github.com/buzbarstow/kosudoku.

### Materials & Correspondence

Correspondence and material requests should be addressed to B.B.. Individual strains (up to ≈ 10 at a time) are available at no charge for academic researchers. We are happy to supply a duplicate of the entire *S. oneidensis* knockout collection to academic researchers, but will require reimbursement for materials, supplies and labor costs. Commercial researchers should contact Cornell Technology Licensing for licensing details.

### Author Contributions

Conceptualization, S.M. and B.B.; Methodology, S.M. and B.B.; Investigation, S.M., A.M.S., B.P. M.R. and B.B; Writing - Original Draft, S.M.; Writing - Review & Editing, S.M., B.B., M.R., M.W., E.G., and B.B.; Funding Acquisition, A.M.S., E.G., M.W., and B.B.; Resources, M.R., E.G., M.W., and B.B.; Supervision, M.R., M.W. and B.B.; Data Curation, S.M. and B.B.; Visualization, S.M. and B.B.; Formal Analysis, S.M..

## Acknowledgements

S.M. was supported by a Cornell Presidential Life Sciences Graduate Fellowship. A.M.S. was supported by a Cornell Energy Systems Institute Postdoctoral Fellowship, and a Small Grant from the Cornell Atkinson Center for Sustainability. This work was supported by Cornell University startup funds, an Academic Venture Fund award from the Atkinson Center for Sustainability at Cornell University, a Career Award at the Scientific Interface from the Burroughs Welcome Fund to B.B., ARPA-E award DE-AR0001341 to B.B, M.H., E.G., and M.W., and a gift from Mary Fernando Conrad and Tony Conrad to B.B..

## Competing Interests

The authors are pursuing patent protection for engineered organisms using knowledge gathered in this work (US provisional patent application number 63/405,353). A.M.S. and S.M. are co-founders (B.B. is a contributor) of REEgen, Inc., which is developing genetically engineered microbes for REE bio-mining.

## Materials and Methods

### Media Preparation

#### bm20 Media

bm20 is composed of 20 mM MES buffer, 8.6 mM ammonium chloride, 0.5 mM magnesium sulfate, 1.7 mM ammonium sulfate, and 5 mL/L Trace Mineral Supplement (ATCC). The media was adjusted to pH 5.5 with 1 M NaOH.

### Genome-wide REE Biosorption Screen

#### Introduction

We screened the *S. oneidensis* whole genome knockout collection [Baym2016a, Anzai2017a] for biosorption of the rare earth element (REE) europium. We hypothesized that using europium, which lies in the middle of the REE size range, would allow us to maximize the number of genes we could discover.

We screened a subset of the knockout collection comprising 77 96-well plates that covers 3,561 unique genes (**Dataset S1**) (the *S. oneidensis* knockout collection is comprised of ≈ 50% blank wells). Due to the failure of some mutants in the collection to grow, we screened 3,373 unique genes in total. Contamination did not appear to be an extensive problem as, on average, we had only 1.6 contaminated wells (wells that were not supposed to have bacteria in them yet grew anyway) per plate. The maximum number of contaminated wells in a plate was 10.

The genetic screen was conducted over the course of several weeks and was divided into batches, each of which took two days to process. Typical batch sizes were between 4 to 8 plates.

#### Replication of *Shewanella oneidensis* Whole-genome Knockout Collection

Microplates from the *S. oneidensis* knockout collection were replicated from a master collection (stored at -80 °Cwith a pin-tool (EnzyScreen CR1000) into a flat-bottom polypropylene plate (Part no. 655261, Greiner) containing 150 μL of LB media per well with 30 mg L^-1^ Kanamycin. Newly inoculated plates were incubated at 30 °C shaking at 800 rpm in a high-throughput microplate shaker (Infors Multitron Pro) for between 16 and 20 hours. The following morning, 3 μL of culture was transferred to a new plate containing fresh LB with 30 mg L^-1^ Kanamycin. The newly diluted cultures were grown for 5-6 hours.

#### Biosorption Assay

Upon removing our plates from the incubator, we diluted 40 μL of culture from each plate into a new round-bottom polystyrene 96-well plate (Part no. 650101, Greiner) with 150 μL of bm20 media (see **Media Preparation**). After this transfer, we took an optical density (OD) measurement of our plate with a plate reader (Biotek Synergy 2) at 590 nm. We called this OD the %growth OD” because it was proportional to the final OD that the bacteria grew to.

After taking the OD, we centrifuged each plate in a swinging bucket centrifuge with micro-well plate adaptors (Eppendorf 5810R) at a speed of 3,214 × *g* for 7 minutes. After that, we positioned a 96-well pipette to remove supernatant from the edges of the wells of the plate. We removed as much supernatant as possible because components of the growth media (data not shown) can bind to rare earth elements and interfere with biosorption. We then rinsed the bacteria by adding 170 μL of bm20 to each well and resuspended by shaking. We then repeated the centrifuging and removing of supernatant steps.

We added 200 μL of bm20 with approximately 25 μM of europium to the rinsed *S. oneidensis* cells and resuspended by vortexing. We then shook the plate in the plate reader or in a plate vortexer for 10 minutes while biosorption occurred. Previous research has shown that bacterial cells tend to reach full biosorption capacity within 10 minutes[Takahashi2005a, Brewer2019]. Upon the completion of the shaking step, we took another OD measurement (called the %final OD”) to have an estimate of how many bacteria were present in each well for the biosorption assay.

We centrifuged each plate once more at a speed of 3214 × *g* for 10 minutes. We then transferred 100 uL of the supernatant to a new flat bottom polystyrene plate (Greiner bio-one ref: 655101). We used the plate reader to take a spectrum of absorbance from 580 nm to 680 nm in increments of 10 nms. These are our %blank” measurements. We then added 100 uL of a solution of 60 μM concentration of Arsenazo III dye dissolved in pH 3.5 20 mM MES buffer to each well with the 96-well pipette. We shook the plate for 4 minutes and then we used the plate reader to measure the absorbance in the same wavelength range as the blanks. These absorbance measurements served as a proxy measurement for the concentration of rare earth elements that were not adsorbed by the bacteria.

### Analysis of Genome-wide REE Biosorption Screen

#### Challenges of Identification of Mutants with Differential Biosorption

Optical density and As-III absorbance measurements were corrected to account for blemishes on the exterior of assay plates, especially due to centrifugation. We knew that it was impossible for our final optical density after rinsing the bacteria to be higher than the optical density prior to rinsing. We thus took our %final” optical density measurement to be the minimum of the optical density prior to rinsing and the optical density after we have finished preparing our biosorption assay. Likewise, As-III absorbance measurements were background corrected by subtracting a blank measurement taken prior to As-III addition.

Non-uniform growth is the biggest challenge in identification of mutants with genetic modifications that affect REE biosorption. Due to the dynamics of ligand-receptor binding chemistry, we would expect that cultures with higher densities will produce higher overall biosorption but have lower biosorption per cell.

We used deviations from a linear piecewise relationship between optical density at 590 nm (OD_590_) and As-III absorbance to identify mutants with truly differential biosorption (**Figure 1B**). The OD to As-III absorbance function varied from plate to plate, likely due to slightly different growth conditions. As a result, we treated every plate separately when identifying outliers.

The linear piecewise function relating OD_590_ and As-III is made of 2 sections. For OD_590_ ⪆ 0.2, As-III absorbance was linearly related to OD_590_. For OD_590_ ⪅ 0.2, we took advantage of the large number of blank wells in each 96-well plate to measure As-III absorbance with no biosorption. Thus, for the earlier part of the piecewise function, we fit a line between a point marking the average OD_590_ and As-III absorbance of blank wells and the point with the smallest OD_590_ in the set used to create the second part of the function. We found that most points in each plate generally fell along our piecewise function.

To find biosorption outliers, we calculated the standard deviation of the distances of each point from the linear piecewise function for each plate. We marked any point as significant if it had a distance of more than 2 standard deviations from our piecewise function (**Figure 1B**).

#### Arsenazo III Assay Quality Control

Quality-control procedures were used to flag low-quality data during analysis. Our procedures specifically looked for anomalous optical density measurements and absorbance spectra patterns.

Plots of final optical density versus growth optical density form a compact line (**Figure S1A**). Datapoints where the final OD_590_ was more than three standard deviations above the average value were flagged. Datapoints where the final OD_590_ was below average were not flagged, as these indicated either that the growth OD_590_ measurement was off (which would not affect the final analysis) or that there was a large loss of bacteria during rinsing (which also would not affect the final analysis).

Arsenazo III absorbance measurements were quality-controlled by ratiometric analysis. Errors in the As-III absorbance measurement can arise due to errors in pipetting the As-III stock, or by the presence of dust or scratches on the assay plate surfaces. For the range of As-III absorbance measurements observed in our assay, the relationship between the 650 nm absorbance and the 680:650 absorbance ratio was linear (**Figures S1B** and **C**). Extreme outliers could occasionally skew this analysis, so we eliminated the five data points furthest from the original linear fit and created a second line of best fit. Data points that lay more than three standard deviations away from the line of best fit were flagged.

Manual inspection was used as the final step in quality control resulting in the selection of 240 mutants for further analysis. Each of the 294 data points that had outlier biosorption measurements in the screen were manually examined. We paid special attention to the datapoints flagged with a high final OD_590_ or with an anomalous absorbance spectrum. We eliminated 12 datapoints that were listed as blank in the *S. oneidensis* knockout collection catalog, but that displayed cross-contamination. We eliminated 7 datapoints that lacked transposon location information. Finally, we removed strains where we judged that the OD_590_ to As-III absorbance piecewise function did not provide a reliable estimate for significance due to lack of surrounding data.

#### Gene Ontology Enrichment Analysis

We followed the gene ontology enrichment analysis procedures laid out in Schmitz *et al*. [Schmitz2021b], except we did not use InterProScan to collect gene ontology data. In brief, we used DIAMOND[Buchfink2021a] to assign annotated protein models with the closest BLAST hit using the Uniref90 database (downloaded from https://www.uniprot.org/uniref/), an E-value threshold of 10^−10^, and a block size of 10. We used the output of this search to assign gene ontologies with BLAST2GO[Gotz2008a].

We performed a gene ontology enrichment analysis with the BioConductor topGO package[Adrian2007a] using the default weight algorithm, the TopGO Fisher test, with a p-value threshold of 0.05.

We performed separate gene ontology enrichment analyses for mutants with significantly higher and lower biosorption (**Figure 1C, Dataset S2**). Following the transposon mutant collection screen, we found that Δ*SO_0625* and Δ*glnA* both produced lower Eu-biosorption and added them to the lower biosorption category.

#### Operon Enrichment Analysis

Operon enrichment analysis was used as a complement to ontology enrichment analysis to identify groups of genes involved in REE-biosorption (**Figure 2, Figure S2**, and **Dataset S3**).

Operon memberships in the *S. oneidensis* genome were predicted by the union of results from operon predictions by MicrobesOnline [Dehal2010a] and ProOpDB [Taboada2012a]. In most cases, the two sources produced highly similar results.

We used Fisher&s exact test to calculate if the set of genes whose disruption conferred differential biosorption was enriched in operons with more than one hit. We compared the total number of gene disruptions that significantly affected biosorption (out of the total number of genes assayed in our genetic screen) to the number of gene disruptions within the operon that significantly affected biosorption (out of the total number of genes we looked at within that operon in our genetic screen). Fisher&s exact test was conducted with the fishertest function in MATLAB.

We also applied Fisher’s exact test to calculate if the sets of gene disruptions that either increased or decreased biosorption were enriched in operons that had more than one gene disruption that increased or decreased biosorption, respectively.

### Confirmation of Transposon Mutant Identity

We validated the identity of transposon mutants from the *S. oneidensis* knockout collection that we conducted follow up analyses on using site specific PCR. The verification reactions used a common primer that bound to the Himar transposon, and a mutant specific primer that bound to the genomic region predicted to be outside the transposon. The identities of all but one transposon mutant was correctly predicted in the *S. oneidensis* knockout collection catalog. The single mutant that was mis-identified was originally annotated as δ*arcA* (δ*SO_3988*), but later found to be 147 bp upstream of *SO_2183*.

### Construction of Gene Deletion Mutants

Clean deletion mutants were constructed to validate the results of transposon screening for *hptA, SO_4685, mshJ*, and *SO_3385* as well as for *glnA* and *SO_0625* which could not be clonally isolated from the knockout collection. Deletions were made by homologous recombination using a suicide vector containing a kanamycin resistance cassette flanked by 1000 bp upstream and downstream sequences surrounding the gene of interest. Mutants that had undergone a second recombination (removing the gene of interest and the kanamycin cassette) or reversion (where the gene of interest was recovered) were selected by a sucrose counter selection. Mutants with a clean deletion were separated from revertants by PCR screening [Rowe2021b].

To ensure that the gene deletion process did not introduce additional changes to the *S. oneidensis* genome, we checked REE-biosorption by revertants recovered in the process of deleting two of the genes. In both cases, REE-biosorption was statistically indistinguishable from the true wild-type (*p*-value < 0.05) (**Figure S5**).

### Analytical Measurement of Biosorption with ICP-MS

We explored biosorption in four different solution conditions (detailed in **Table 1**) with three rare earth elements: La (representing light REE), Eu (representing middle REE), and Yb (representing heavy REE). In every condition, bacterial culture density was normalized to the same optical density.

Bacterial strains of interest were retrieved from glycerol stocks frozen at -80 °C and recovered on LB agar plates (with 50 mg L^-1^ kanamycin for transposon insertion strains). We picked three single colonies for each strain and inoculated them into separate wells containing 200 μL of LB (with 50 mg L^-1^ kanamycin for transposon insertion strains) in 96-well flat-bottom polypropylene plates (Greiner Bio-One ref: 655261) and incubated them at 30 °C overnight shaking at 800 rpm.

The following morning, we back-diluted 30 μL from each well into culture tubes containing 3 mL of LB (with 50 mg L^-1^ kanamycin for transposon mutants). Cultures were incubated at 30 °C until they reached an optical density (OD_590_) of between 1.3 and 1.45.

Each culture was used for 4 biosorption experiments in each of the different conditions detailed in **Table 1**. Each culture was split into two 1.7 mL centrifuge tubes, pelleted at 7,800 × *g*, resuspended in 1 mL of buffer, and then pelleted one more time at 7,800 × *g*. The first tube was rinsed with a low ionic strength buffer (20 mM MES, 20 mM NaCl, adjusted to pH 5.5 with 5M NaOH) and the second tube with a high ionic strength buffer (20 mM MES, 100 mM NaCl, adjusted to pH 5.5 with 5M NaOH).

We resuspended the rinsed cells in 600 μL of the same respective buffer, took the optical density (OD), then divided each culture into two new tubes with a final OD of 0.85 and either high or low REE concentrations as follows. For cultures in low ionic strength solution (final concentrations of 10 mM NaCl and 10 mM MES), the low REE solution contained 30 μM each of lanthanum, europium, and ytterbium and the high REE solution contained 60 μM of each. For cultures in high ionic strength solution (final concentrations of 50 mM NaCl and 10 mM MES), the low REE solution contained 10 μM each of lanthanum, europium, and ytterbium and the high REE solution contained 30 μM of each.

### ICP-MS Measurements

Our ICP-MS samples were prepared by diluting our biosorption samples 1/25 in 2% trace metal grade nitric acid (JT9368, J.T. Baker, Radnor, PA). Our samples were analyzed using an Agilent 7800 ICP-MS (m/z: La, 139; Eu, 151; Yb, 172) using a rare earth element mix standard which included all the other rare earth elements in addition to the three we analyzed (67349, Sigma-Aldrich, St. Louis, MO) and a rhodium in-line internal standard (SKU04736, Sigma-Aldrich, St. Louis, MO, m/z = 103). ICP-MS data were analyzed using the program MassHunter, version 4.5. Quality control was conducted by doing periodic measurements (every ten samples) of our standards (the 10, 25, 50, and 100 ppb) and 2% nitric acid blanks. We used the Rh internal standard to account for effects of drift. Repeat standards were analyzed periodically (or every 10 samples) and were quantified with an accuracy of +/- 2.5%

### Comparing Transposon Containing and Wild-Type S. oneidensis Strains

We found that the average biosorption of transposon insertion strains did not resemble our wild-type bacteria. We theorize that even if the gene a transposon was inserted into did not alter biosorption, it is possible that the transposon itself—or the fact that the insertion mutant strains were grown up with Kanamycin—impacts biosorption. To test this hypothesis, we took four transposon mutants (which we refer to as quasi-wild-type or qWT) that had the transposon in a presumably neutral location and whose biosorption in the As-III screen did not significantly differ from average mutant in the containing plate.

### Choice of Quasi-WT Strains

We expected that a transposon appearing at the end of a gene would have no effect on that gene. We thus picked transposon mutants where the transposon appeared at the very end of the gene. We also ensured that the transposon was at least 300 base pairs away from the start of any other gene to minimize disruption of promoter regions. Finally, we confirmed that the selected disruptions did not have any significant changes in biosorption within our assay. Four transposon mutants were selected at random from the mutants that met these requirements. The end of the genes where the transposon appeared were *SO_4279* for qWT_1_, *SO_4707* for qWT_2_, *SO_0214* for qWT_3_, and *SO_2225* for qWT_4_.

Biosorption of each qWT was compared to that of the natural WT and the other qWT strains using a two-sided t-test. We found that the wild-type showed at least 13% higher biosorption compared to the average qWT in every solution condition (**Figure S4**). Additionally, the qWT had solution condition-dependent differences in total biosorption compared to each other. qWT_4_ had higher (*p* < .05) biosorption than the average qWT for the HH solution condition. When we performed pairwise t-tests between our four qWT mutants, we found that in three out of four of our solution conditions, we had some mutant or mutants that had different biosorption than the others. In LL, qWT_2_ had significantly higher biosorption than each of the other qWTs. In LH, qWT_2_ had higher biosorption than qWT_3_ and qWT_4_. In HH, qWT_4_ had significantly higher biosorption than every other qWT and qWT_3_ had significantly lower biosorption than qWT_1_.

### Methodology for Comparing Relative REE Biosorption

We compared the amount of biosorption for each individual REE to the total REE biosorption for each solution condition. We found that, over a finite range of total REE biosorption, there was generally a linear relationship between individual and total REE biosorption. We thus used our data for our transposon insertion mutants to plot lines of best fit in each solution condition comparing each individual REE biosorption to total REE biosorption. We speculated that, even though our data set consisted almost totally of strains that had outlier biosorption compared to the wild-type, the relative biosorption changes would not be in any particular direction. We thus expect the baseline calculated from these mutants to reasonably resemble the true average of *S. oneidensis* transposon insertion strains.

We excluded our two biggest disruption mutant total biosorption outliers—δ*nusA* and δ*SO_4685*—from our analysis. We left these strains out of our analysis because their total REE biosorption fell outside of the finite range of total biosorption that we felt confident was linear with individual REE biosorption.

Once we had our line of best fit, *f* (*REE*_*T*_), for each data point, we calculated the percent change of biosorption of the individual REE of interest (REE_i_) on the y-axis compared to the expected value based on the total REE biosorption (*REE*_*T*_) on the x-axis: 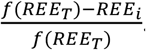. For each strain we calculated the mean and standard error of this percent change. Significance was calculated by doing a two-sided t-test looking to see if our percent change of biosorption was significantly different than 0.

#### Effects of Extra Incubation Time on Biosorption

Previous research [Takahashi2005a] has found that, in a sufficiently high pH environment, the longer bacteria were exposed to REE, the less biosorption occurred. The authors theorized that this decrease in biosorption over time was caused by bacterial secretions. Since they found that this occurred at pH 5.8 and we conducted our assays at pH 5.5, we conducted experiments to test how much different incubation times could have affected our results. Since the amount of time we let our bacteria mix with REE was fixed inside our experiment, we chose instead to conduct our experiments by changing the amount of time we let our bacteria sit after rinsing was completed, but prior to adding the REE for our assays.

We tested the effects of secretions in two of our biosorption environments, LL and HH. We did not find a statistically significant impact on absolute biosorption or on the separation factor for LL. While our results were not statistically significant, it did appear like there was a clear downward trajectory to the level of biosorption as well as an increase in the Yb/La and Eu/La separation factors. For HH, on the other hand, there was a substantial decrease in the overall biosorption level as well as a substantial increase in the Yb/La and Eu/La separation factors. However, the overall biosorption decrease between the 74 minute and 138 minute waiting periods lacked statistical significance and was far less than the decrease between the 42 and 74 minute measurements. This suggests that the effects of the secretions (or whatever other mechanism is responsible for the change in REE binding) decrease over time—a result that would give us confidence in our experiments given that our bacteria typically sat in their final assay solutions for between 90 and 120 minutes prior to the completion of our biosorption assay.

### Statistical Information

Statistics relating to the genetic screen (identifying outliers, gene ontology enrichment analysis, operon enrichment analysis) are all described in their respective sections. All other statistics were performed using two-tailed t-tests with three biological replicates each.

