## Supplementary Information for "Genomic Characterization of Rare Earth Binding by *Shewanella oneidensis*"

[†]Corresponding author:

### Supplementary Information Figures

**Figure S1.** Absorbance measurements were used to perform quality-control tests on the Arsenazo III screen for differential Eu-biosorption.

**Figure S2.** Thirteen operons are significantly enriched in genes influencing biosorption.

**Figure S3.** REE-biosorption and separation factor appear to equilibrate during the incubation time used in measurements in this study.

**Figure S4.** REE-biosorption by quasi-wild-type strains of *S. oneidensis* is lower than the true wild-type.

**Figure S5.** Recombined wild-type *S. oneidensis* strains do not have significantly different biosorption compared to the original wild-type.

**Figure S6.** ICP-MS measurements find 23 transposon insertion mutants of *S. oneidensis* with statistically significant changes in relative REE-biosorption in at least one solution environment, although few of these changes are robust.

### Supplementary Information Tables

**Table S1.** 29 Genes that control Eu-biosorption are also regulated by the Arc system.

**Table S2.** ICP-MS measurements validate the results of high-throughput Eu-biosorption screening in up to 79% of cases.

### Supplementary Information Datasets

**Dataset S1.** The high-throughput Arsenazo III screen found 242 genes that produced differential Eu-biosorption.

**Dataset S2.** The high-throughput Arsenazo III screen identified 18 gene ontologies involved in Eu-biosorption.

**Dataset S3.** Operon enrichment analysis found thirteen operons involved in Eu-biosorption.

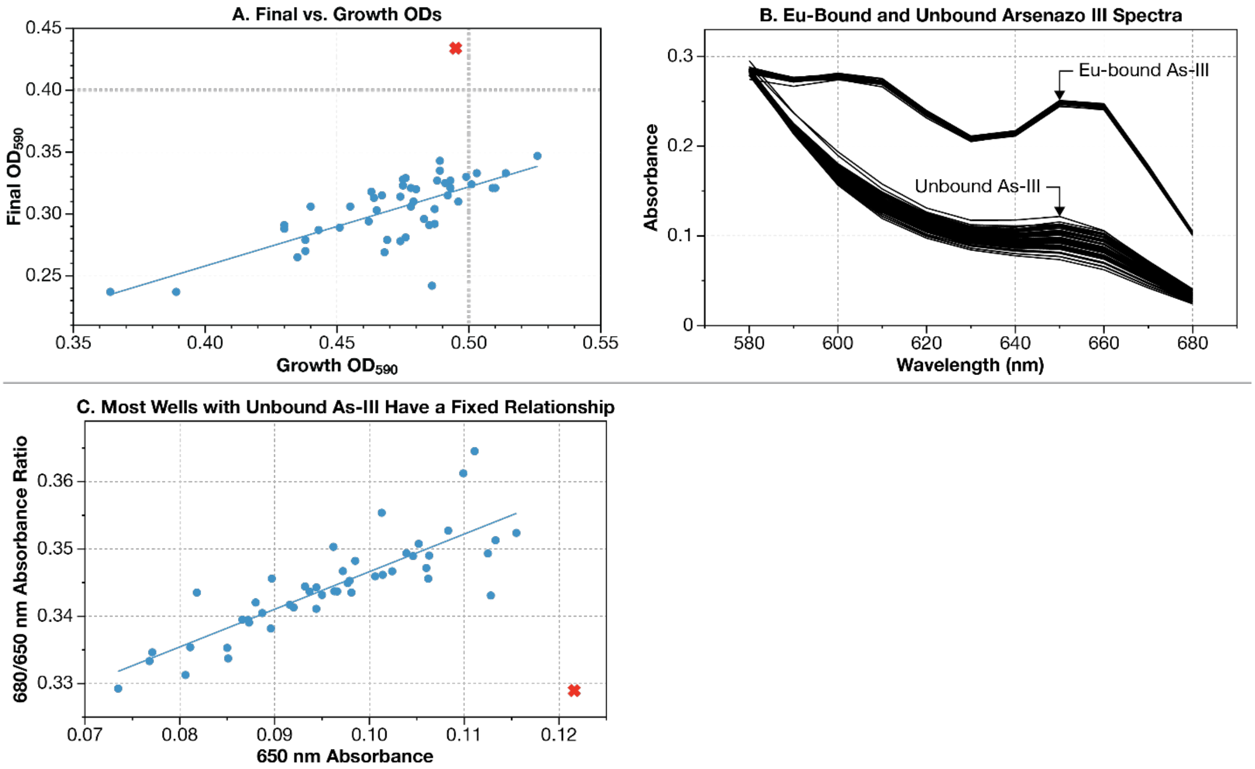

**Figure S1. Quality control for the Arsenazo III screen for differential Eu-biosorption.** Complement to **Figure 1** in the main text. (**A**) Prior to the high-throughput Arsenazo III assay, mutants from the *S. oneidensis* whole genome knockout collection [Baym2016a, Anzai2017a] were grown to saturation in 96-well plates, and their optical density was measured with a plate reader (indicated on the *x*-axis as Growth $OD_{590}$). We next rinsed the bacteria in our 96-well plate, mixed the bacteria with REE, and re-measured the optical density (indicated on the y-axis as Final $OD_{590}$). We observed a linear relationship between Growth $OD_{590}$ and Final $OD_{590}$ for the vast majority of wells (blue circles). Wells that had a significantly higher Final $OD_{590}$ than expected (the single red cross) were flagged for manual inspection. (**B**) The Arsenazo III dye shows an increase in absorbance at ≈ 650 nm when bound to Eu. (**C**) Almost all wells tested in the Arsenazo III assay show a linear relationship between absorbance at 650 nm and the ratio of absorbances at 680 and 650 nm (blue circles). Wells that significantly deviated from this relationship (the single red cross) were flagged for manual inspection.

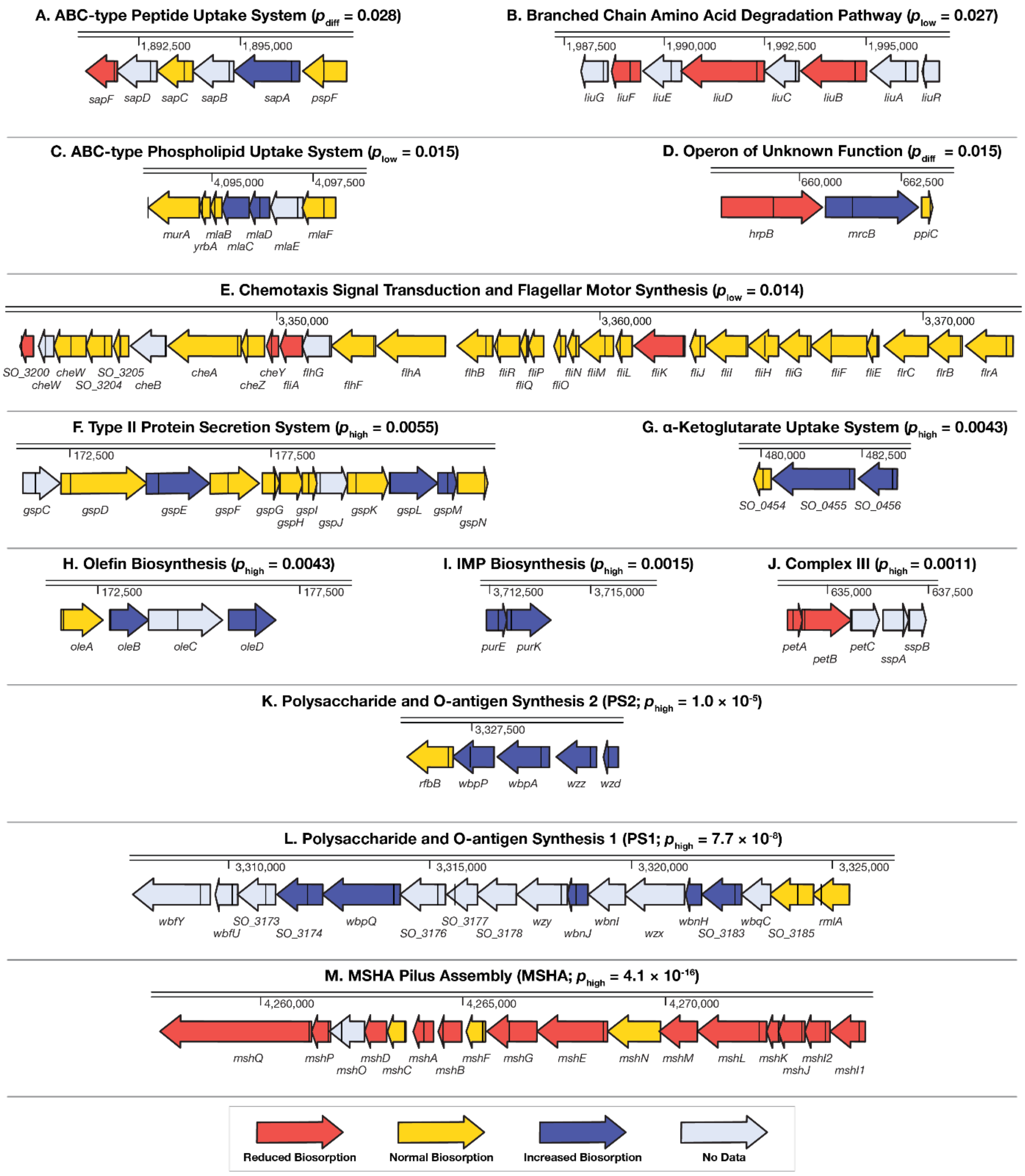

**Figure S2. Thirteen operons are significantly enriched in genes influencing REE biosorption.** Complement to **Figure 2**. The results of the high-throughput Eu-biosorption screen (**Dataset S1**) of the *S. oneidensis* knockout collection [Baym2016a, Anzai2017a] were analyzed to find operons with statistically-significant enrichments of hits (**Dataset S3**). The location of the transposon disruption in each gene (found in the *S. oneidensis* whole genome knockout collection catalog [Baym2016a]) is marked as a black line.

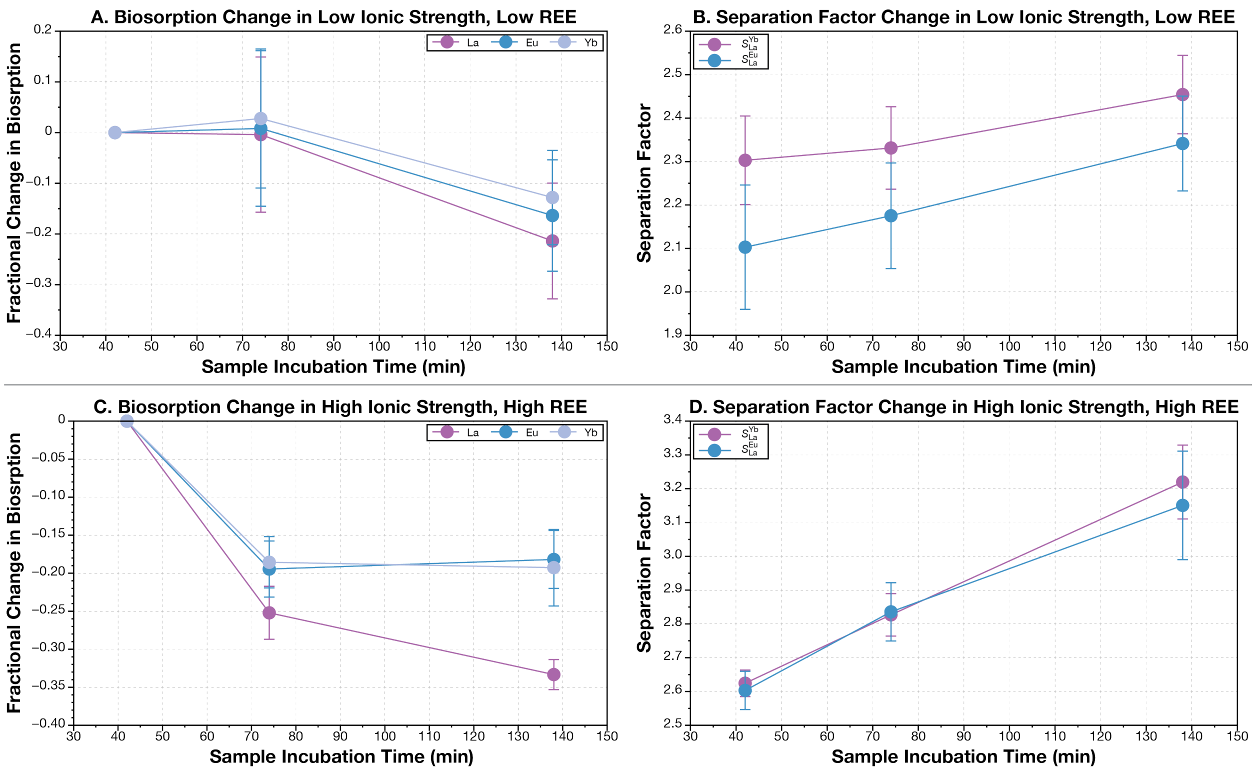

**Figure S3. Effects of extra incubation time on REE-biosorption and separation factor appear to be small.** We tested the effects of incubation time in two of our biosorption environments: low ionic strength, low total REE (LL) and high ionic strength, high total REE (HH) (see **Table 1** for full details). (**A** and **B**) We did not find a statistically significant impact on absolute biosorption or on the separation factor with increasing incubation time under LL. While our results were not statistically significant, it did appear like there was a clear downward trajectory to the level of biosorption as well as an increase in the Yb/La and Eu/La separation factors. (**C** and **D**) For HH, there was a statistically significant decrease in the overall biosorption level as well as a significant increase in the Yb/La and Eu/La separation factors. There was not, however, a significant difference in overall biosorption level between 74 and 138 minute incubation times. Considering that incubation times for our actual assay were generally between 90 and 120 minutes, we conclude that differences in incubation time likely did not have a large impact on our results. Error bars indicate standard deviation of three biological replicates.

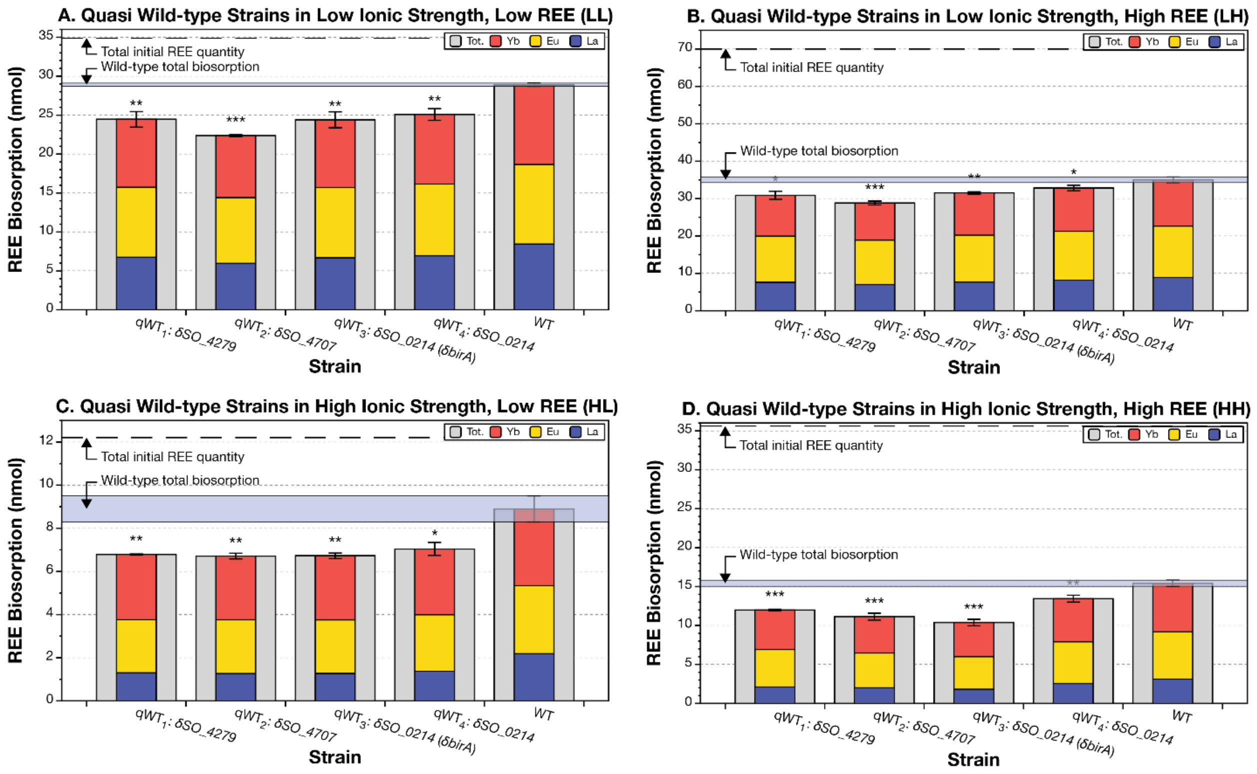

**Figure S4. REE-biosorption by quasi-wild-type strains of *S. oneidensis* are different from each other and are all lower than the true wild-type.** We selected 4 'quasi-wild-type' (qWT) transposon insertion mutants from the *S. oneidensis* whole genome knockout collection [Baym2016a, Anzai2017a]. These qWT mutants did not cause any differential biosorption in the As-III Eu-biosorption screen and featured a transposon insertion towards the end of a protein coding region where it was least likely to impact function. We measured REE biosorption for each of these strains under four solution conditions shown in panels **A** to **D** (solution conditions are detailed in **Table 1)**. As biosorption by qWT strains and wild-type (WT) are different, we used the average total and individual REE-biosorption by the qWT mutants for comparison with selected mutants of interest in **Figures 3**, **4**, and **S4**. (**A**) Under the low ionic strength, low REE environment, $qWT_2$ has significantly lower biosorption than $qWT_3$ and $qWT_4$. (**B**) Under the low ionic strength, high REE environment, $qWT_2$ has significantly lower biosorption than all the other qWTs. (**C**) Under the high ionic strength, low rare earth environment, none of the qWTs are significantly different from each other. (**D**) Under the high ionic strength, high REE environment, $qWT_4$ has significantly higher biosorption than all the other qWTs and $qWT_3$ has significantly lower biosorption than $qWT_1$. The number of stars above or below each bar indicates the statistical significance of the measurement difference from wild-type: *: $p$-value $< 0.05$; **: $p$-value $< 0.01$; ***: $p$-value $< 0.001$. Error bars show standard deviation of three biological replicates.

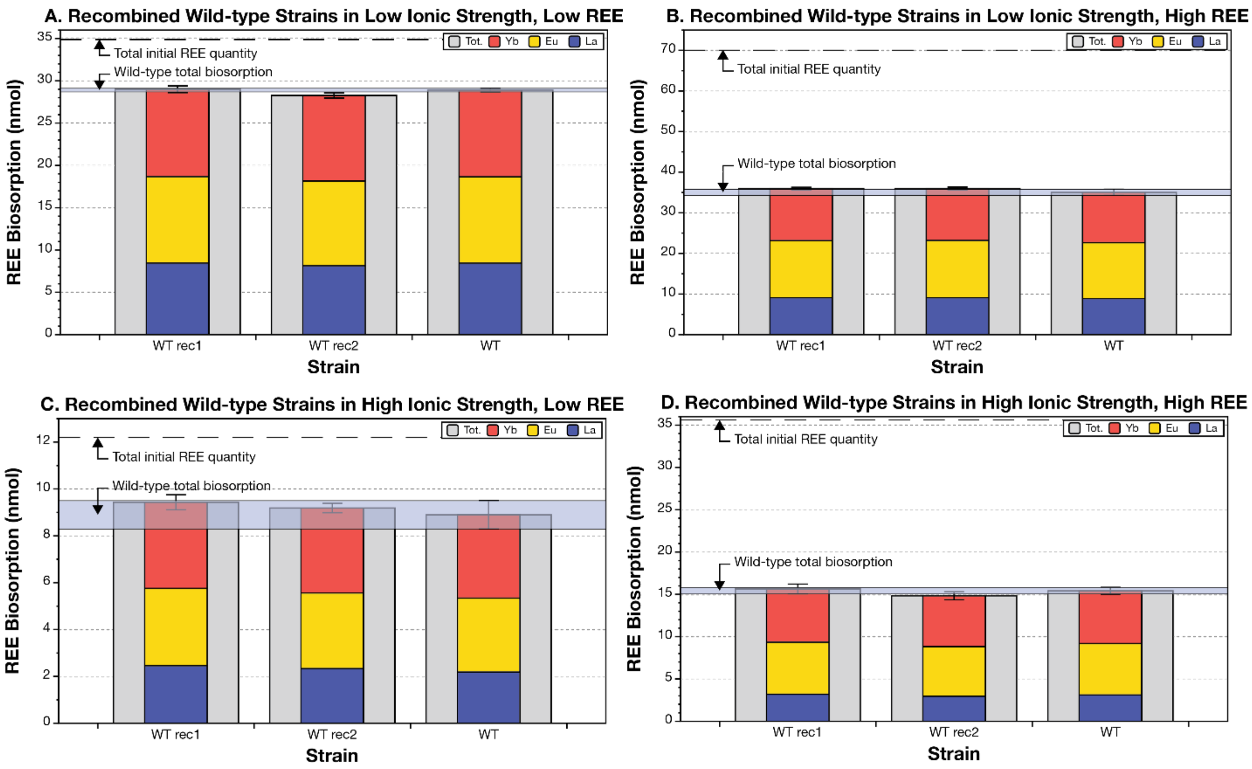

**Figure S5. Recombined wild-type *S. oneidensis* strains do not have significantly different biosorption compared to the original wild-type.** We used two homologous recombination steps in order to delete our gene of interest. After the second step, we could end up with a gene deletion genotype or with the wild-type. To confirm that the process of deleting the gene of interest did not cause any changes in biosorption, we tested the biosorption of some of our recombinants that re-created the wild-type genome. As expected, we found that there was no significant alteration in biosorption of our recombinants compared to our original wild-type. The number of stars above or below each bar indicates the statistical significance of the measurement difference from wild-type: *: $p$-value < 0.05; **: $p$-value < 0.01; ***: $p$-value < 0.001. Error bars show standard deviation of three biological replicates.

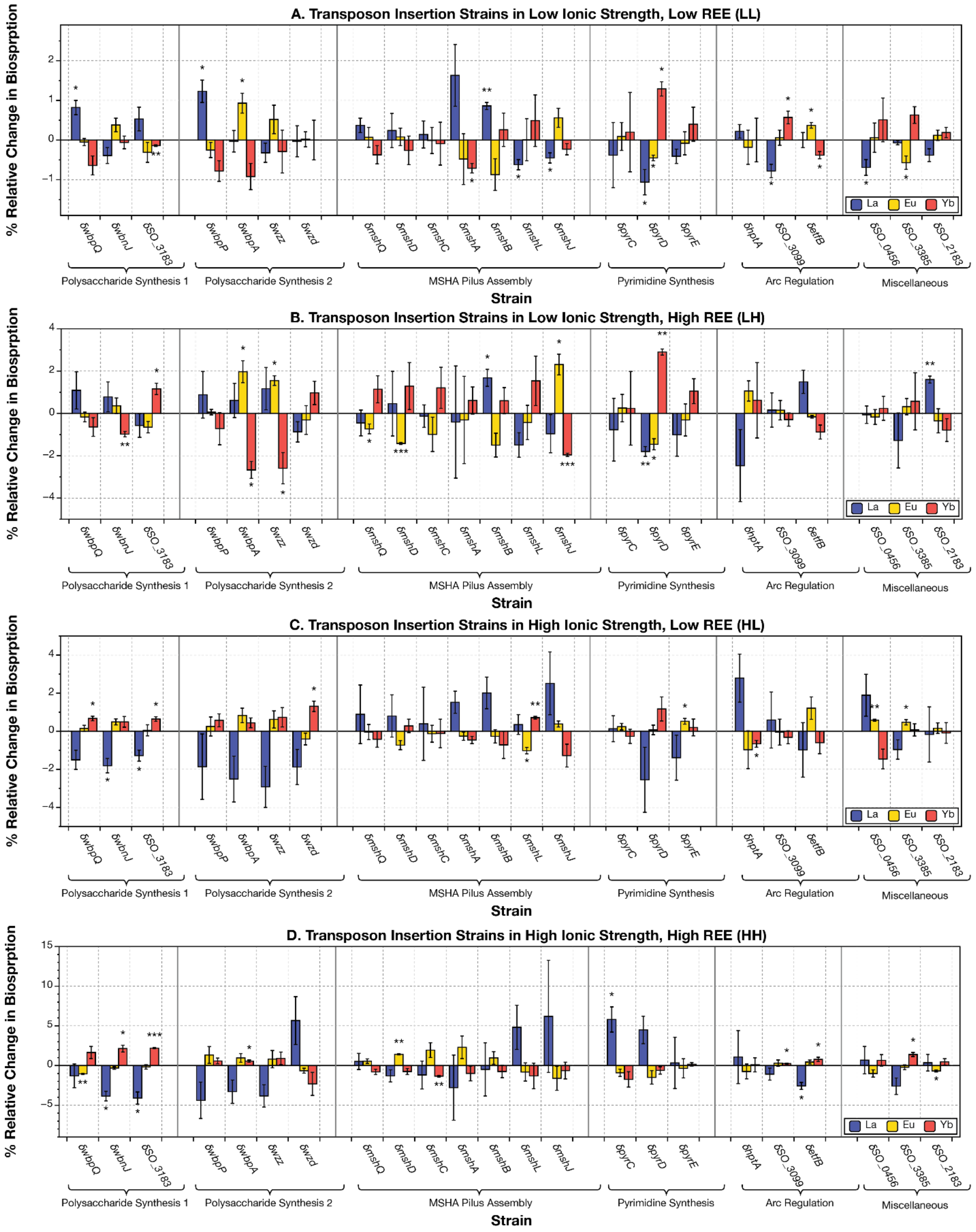

**Figure S6. ICP-MS measurements find all transposon insertion mutants of *S. oneidensis* tested register statistically significant changes in relative REE-biosorption in at least one solution environment, although few of these changes are robust.** This figure is a complement to **Figure 4** in the main text. The number of stars above or below each bar indicates the statistical significance of the measurement difference from quasi-wild-type: *: $p$-value < 0.05; **: $p$-value < 0.01; ***: $p$-value < 0.001. δ indicates a transposon insertion mutant. Error bars show standard deviation of three biological replicates. Details of solution conditions shown in panels (**A**) to **(D**) can be found in **Table 1** in the main text.

| ORF | Gene | log(*arcS* Deletion Activity/WT Activity) | Arc Regulatory Mode | Knockout Biosorption Relative to Wild-type | Gene Effect on Biosorption | Scenario |
|---|---|---|---|---|---|---|
| SO_2091 | *hypD* | 3.3 | ⊣ | Lower | ↑ | 1 |
| SO_4157 | *ttrR* | 2.07 | ⊣ | Lower | ↑ | 1 |
| SO_2493 | *psrA* | 1.34 | ⊣ | Lower | ↑ | 1 |
| SO_3145 | *etfB* | 1.33 | ⊣ | Lower | ↑ | 1 |
| SO_4266 | *fic* | 1.32 | ⊣ | Lower | ↑ | 1 |
| SO_0696 | *dsbD* | 1.2 | ⊣ | Lower | ↑ | 1 |
| SO_3774 | *putA* | 1.17 | ⊣ | Lower | ↑ | 1 |
| SO_0261 | *ccmC* | 1.02 | ⊣ | Lower | ↑ | 1 |
| SO_3967 | *SO_3967* | -1.08 | → | Higher | ↓ | 2 |
| SO_3877 | *SO_3877* | -1.09 | → | Higher | ↓ | 2 |
| SO_4255 | *pyrE* | -1.16 | → | Higher | ↓ | 2 |
| SO_0761 | *glnK* | -1.35 | → | Higher | ↓ | 2 |
| SO_3099 | *SO_3099* | -6.4 | → | Higher | ↓ | 2 |
| SO_0090 | *SO_0090* | 3.47 | ⊣ | Higher | ↓ | 3 |
| SO_3422 | *yfiA* | 2.2 | ⊣ | Higher | ↓ | 3 |
| SO_1483 | *aceB* | 1.83 | ⊣ | Higher | ↓ | 3 |
| SO_4730 | *hemN* | 1.53 | ⊣ | Higher | ↓ | 3 |
| SO_3980 | *nrfA* | 1.43 | ⊣ | Higher | ↓ | 3 |
| SO_1343 | *rseA* | 1.39 | ⊣ | Higher | ↓ | 3 |
| SO_0174 | *gspL* | 1.2 | ⊣ | Higher | ↓ | 3 |
| SO_0839 | *SO_0839* | -1.03 | → | Lower | ↑ | 4 |
| SO_1167 | *mrdB/rodA* | -1.04 | → | Lower | ↑ | 4 |
| SO_0468 | *ubiA* | -1.09 | → | Lower | ↑ | 4 |
| SO_4069 | *SO_4069* | -1.2 | → | Lower | ↑ | 4 |
| SO_2755 | *rnt* | -1.38 | → | Lower | ↑ | 4 |
| SO_2111 | *SO_2111* | -1.44 | → | Lower | ↑ | 4 |
| SO_4178 | *mxdC* | -1.58 | → | Lower | ↑ | 4 |
| SO_0095 | *hutI* | -1.64 | → | Lower | ↑ | 4 |
| SO_2389 | *emrD* | -3.55 | → | Lower | ↑ | 4 |

**Table S1. 29 Genes that control Eu-biosorption are also regulated by the Arc system.** We found 29 genes whose disruption changes Eu-biosorption amongst the 604 genes controlled by the Arc system [Lassak2013a]. If gene expression is increased by deletion of *arcS* (measured in Lassak *et al.* [Lassak2013a]), this suggests that the gene is repressed (⊣) by the Arc system. Alternatively, if expression goes down, the gene is activated (→) by Arc. If gene knockout reduces biosorption relative to wild-type, then the gene promotes REE biosorption (↑). Likewise, if the knockout increases biosorption, then the gene discourages it (↓). We speculate that all of these effects are relative to a moderately hypoxic environment since liquid in microwell plates used in high-throughput screening is typically not well mixed. We envision four scenarios in which disruption of the Arc system (by disruption of *hptA*) can affect biosorption: (1) de-repression of genes that promote REE biosorption; (2) a failure of activation of genes that discourage REE biosorption; (3) de-repression of genes that discourage REE biosorption and (4) failure of activation of genes that promote biosorption. The increase in biosorption by disruption of hptA suggests to us that the combined effects of scenarios 1 and 2 dominate the combined effects of scenarios 3 and 4.

| Locus | Gene Disruption Mutant | Protein Product | Gene Group | As-III | LL | LH | HL | HH |
|---|---|---|---|---|---|---|---|---|
| SO_3189 | *wbpP* | UDP-GlkcNAc C4 epimerase WbpP | PS1 | H | H | H | NSC | NSC |
| SO_3190 | *wbpA* | UDP-N-acetyl-d-glucosamine 6-dehydrogenase WbpA | PS1 | H | H | H | NSC | NSC |
| SO_3191 | *wzz* | polysaccharide chain length determinant Wzz | PS1 | H | NSC | NSC | NSC | NSC |
| SO_3192 | *wzd* | hypothetical protein | PS1 | H | H | H | H | H |
| SO_3175 | *wbpQ* | asparagine synthase glutamine-hydrolyzing WbpQ | PS2 | H | H | H | H | H |
| SO_3183 | *SO_3183* | perosamine synthetase-related protein | PS2 | H | H | H | H | H |
| SO_4799 | *wbnJ* | O-antigen biosynthesis acetyltransferase WbnJ | PS2 | H | H | H | H | H |
| SO_4100 | *mshQ* | MSHA pili-associated adhesin MshQ | MSHA | L | NSC | NSC | H | H |
| SO_4103 | *mshD* | MSHA minor pilin protein MshD | MSHA | L | NSC | NSC | NSC | H |
| SO_4104 | *mshC* | MSHA minor pilin protein MshC | MSHA | NSC | NSC | NSC | H | H |
| SO_4105 | *mshA* | MSHA major pilin subunit MshA | MSHA | L | NSC | L | NSC | NSC |
| SO_4106 | *mshB* | MSHA minor pilin protein MshB | MSHA | L | NSC | L | H | NSC |
| SO_4112 | *mshL* | MSHA system outer membrane secretin MshL | MSHA | L | H | NSC | NSC | NSC |
| SO_4114 | *mshJ* | MSHA biogenesis protein MshJ | MSHA | L | NSC | NSC | L | L |
| SO_2592 | *pyrD* | dihydroorotate dehydrogenase PyrD | Pyrimidine | H | NSC | NSC | H | NSC |
| SO_3695 | *pyrC* | dihydroorotase homodimeric type PyrC | Pyrimidine | H | NSC | NSC | H | NSC |
| SO_4255 | *pyrE* | orotate phosphoribosyltransferase PyrE | Pyrimidine | H | NSC | NSC | H | NSC |
| SO_1327 | *hptA* | histidine-containing phosphotransfer domain protein HptA | Arc | H | H | H | H | H |
| SO_3099 | *SO_3099* | outer membrane long-chain fatty acid receptor FadL family | Arc | H | H | H | H | H |
| SO_3145 | *etfB* | electron transfer flavoprotein beta subunit EtfB | Arc | L | H | H | H | H |
| SO_0456 | *SO_0456* | alpha-ketoglutarate uptake system substrate-binding component | Diverse | H | H | H | H | NSC |
| SO_1203 | *nusA* | N utilization substance protein A NusA | Diverse | L | L | L | L | L |
| SO_2183 | *SO_2183* | LD-transpeptidase ErfK/YbiS/YcfS/YnhG family | Diverse | H | H | H | H | H |
| SO_3385 | *SO_3385* | transcriptional activator of singlet oxygen protection | Diverse | H | H | H | H | NSC |
| SO_4685 | *SO_4685* | outer membrane protein in capsule/EPS biosynthesis locus | Diverse | H | H | H | H | H |
| | | | | ***Number of significant increases or decreases in total biosorption*** | 15 | 16 | 19 | 14 |
| | | | | ***Number of matches to As-III screen*** | 13 | 15 | 15 | 10 |
| | | | | ***Number of matches to As-III screen under any condition*** | 19 | | | |

**Table S2. ICP-MS measurements validate the results of high-throughput Eu-biosorption screening in up to 79% of cases.** This table is a complement to **Figure 3** in the main text. Twenty-five genes highlighted by high-throughput screening with the Arsenazo-III (As-III) assay (**Dataset S1**) were selected for further analysis by mass spectrometry in four solution conditions (detailed in **Table 1** in the main text):

low ionic strength, low total initial REE (LL); low ionic strength, high total initial REE (LH); high ionic strength, low total initial REE (HL); and high ionic strength, high total initial REE (HH). H: higher biosorption than quasi-wild-type; L: lower total lanthanide biosorption than quasi-wild-type; NSC: no significant change. PS1: Polysaccharide and O-antigen Synthesis Operon 1; PS2: Polysaccharide and O-antigen Synthesis Operon 2.
